# ER-vacuole tether Mdm1 coordinates lipid homeostasis to support autophagosome formation

**DOI:** 10.64898/2026.11.18.766846

**Authors:** Eseiwi Obaseki, Daniel Adebayo, Mohamad Rahal, Gonçalo Vale, Jeffrey G. McDonald, Hanaa Hariri

**Affiliations:** Biological Sciences Department, Wayne State University, Detroit, MI, USA; Center for Human Nutrition, UTSW Medical Center, Dallas, Texas, USA; Department of Internal Medicine, UTSW Medical Center, Dallas, Texas, USA; Department of Molecular Genetics, UTSW Medical Center, Dallas, Texas, USA; Department of Oncology, Wayne State University School of Medicine, Detroit, MI, USA

**Author notes:** **Correspondence:** Hanaa Hariri, PhD.

## Abstract

Autophagosome formation requires cells to mobilize lipids for phagophore expansion and completion while maintaining lipid homeostasis during nutrient stress. How lipid intermediates are partitioned between membrane synthesis and neutral lipid storage under these conditions remains unclear. Here, we identify ER-vacuole tether Mdm1 as a regulator of starvation-induced autophagy in budding yeast. Loss of Mdm1 reduced autophagic flux despite preserved TORC1 responses to starvation and impaired autophagosome completion, as indicated by the accumulation of perivacuolar Atg8-positive structures and reduced protease protection. Lipidomic analyses revealed decreased phosphatidylinositol (PI) in whole cells and Atg8-enriched membrane fractions, accompanied by increased diacylglycerol (DAG) and storage lipids. Derepression of phospholipid synthesis through OPI1 deletion or inhibition of triacylglycerol synthesis restored autophagic flux. Remarkably, exogenous DAG also rescued autophagy despite elevated endogenous DAG in Mdm1-deficient cells. DAG supplementation increased PI abundance and improved protease protection, linking changes in lipid metabolism to autophagosome completion. Together, these findings implicate Mdm1 in coordinating membrane phospholipid homeostasis with neutral lipid metabolism and suggest that the metabolic availability of lipid intermediates contributes to efficient autophagosome formation during starvation.

**Summary:** The ER-vacuole tether Mdm1 supports autophagosome completion during starvation by maintaining lipid homeostasis. Loss of Mdm1 reduces phosphatidylinositol and increases neutral lipid accumulation, while genetic or metabolic interventions that alter lipid synthesis restore autophagic flux. These findings link the balance between membrane and storage lipids to efficient autophagosome formation.

## Introduction

Autophagy allows cells to survive nutrient limitation by delivering cytoplasmic material to the lysosome or, in yeast, the vacuole for degradation and recycling. During starvation, this process depends on the rapid formation of autophagosomes, double-membrane vesicles that form when a precursor membrane, the phagophore, expands around cytoplasmic cargo and seals before fusing with the degradative compartment (Reggiori & Klionsky, 2013). Within minutes of induction, the phagophore must acquire sufficient lipids of the appropriate composition to expand and complete autophagosome formation.

Much of these lipids are synthesized on demand. The acyl-CoA synthetase Faa1 accumulates on nucleated phagophores and channels fatty acids into *de novo* phospholipid synthesis in the adjacent endoplasmic reticulum (ER) (Schütter et al., 2020a), while the lipid transfer proteins Atg2 and Vps13 transport lipids from the ER to the expanding phagophore (Dabrowski et al., 2023; Osawa et al., 2019; Valverde et al., 2019). Lipid composition is also important for autophagosome formation. Phosphatidylinositol (PI) is a major component of yeast autophagic membranes (Schütter et al., 2020a) and is abundant in autophagic membranes in *Drosophila (Laczkó-Dobos et al., 2021)*. In mammalian cells, autophagosomes are initiated at ER subdomains enriched in PI synthase (Nishimura et al., 2017). PI also serves as the precursor of the phosphoinositides, including PI 3-phosphate (PI3P) and PI 4-phosphate (PI4P), which regulate autophagy machinery and membrane dynamics (Baba & Balla, 2020; Kurokawa et al., 2019). Perturbations in phospholipid composition can impair autophagosome completion (Polyansky et al., 2022).

The precursors required for membrane phospholipid synthesis are also used for neutral lipid storage. Phosphatidic acid (PA) is a key metabolic branch point: Pah1 dephosphorylates PA to diacylglycerol (DAG), which can be converted to triacylglycerol (TAG) for storage, whereas Dgk1 phosphorylates DAG back to PA, providing a route to phospholipid synthesis (Carman & Han, 2006; Han et al., 2006; Henry et al., 2012). Starvation activates Pah1 through the Nem1-Spo7 phosphatase, and this activation is required for autophagy (Rahman et al., 2018). Neutral lipid synthesis also contributes to autophagy in starved yeast (Velázquez et al., 2016)(Li et al., 2015; Shpilka et al., 2015). However, blocking TAG synthesis can cause DAG accumulates in the ER and disrupt endomembrane organization (Li et al., 2020). Moreover, DAG exists in pools that differ in location and metabolic fate (Ganesan et al., 2015) (Eichmann & Lass, 2015). Thus, pathways supporting lipid storage and membrane synthesis are interconnected, but how their activities are coordinated to meet the lipid demands of autophagosome formation remains unclear.

Membrane contact sites are well positioned to coordinate lipid metabolism because they facilitate lipid exchange between organelles and concentrate lipid-metabolic enzymes in defined locations (Reinisch et al., 2025). In budding yeast, the nucleus-vacuole junction (NVJ), a contact site between the nuclear ER and the vacuole, is enriched in lipid-metabolic enzymes and expands under nutrient stress (Bandyopadhyay et al., 2024; Hariri et al., 2018). The properties suggest that the NVJ may help coordinate lipid metabolism during adaptation to starvation.

Mdm1 is an ER-anchored tether at the NVJ that binds the vacuole through its phosphoinositide-binding phox homology (PX) domain (Henne et al., 2015). During nutrient stress, Mdm1 binds fatty acids, cooperates with Faa1, and defines sites of lipid droplet (LD) formation adjacent to the vacuole (Hariri et al., 2018, 2019). Mdm1 belongs to the sorting nexin-regulator of G-protein signalling (SNX-RGS) family, whose metazoan members SNX13, SNX14 and SNX25 have been linked to autophagy (Akizu et al., 2015; Lauzier et al., 2022; Zheng et al., 2006). Structural studies have identified a conserved hydrophobic cavity formed by the PX-associated A and C (PXA and PXC) domains (Aboumourad & Hariri, 2026; Paul et al., 2022). Nvj3, a yeast paralogue of Mdm1 containing of these two domains, binds PA (Gov et al., 2026) and regulates Dga1-mediated TAG synthesis (Adebayo, Obaseki, Miller, et al., 2026). Together, these observations suggest that SNX-RGS proteins participate in lipid homeostasis, but whether Mdm1 also contributes to autophagosome formation remains unknown.

Here, we show that Mdm1 supports starvation-induced autophagy in *Saccharomyces cerevisiae*. Loss of Mdm1 reduces autophagic flux despite perserved target of rapamycin complex 1 (TORC1) responses to starvation and is associated with impaired autophagosome completion. During nitrogen starvation, Mdm1-deficient cells accumulate DAG, TAG, and LDs, while PI is reduced in whole cells and Atg8-enriched membrane fractions. Derepressing phospholipid synthesis through OPI1 deletion or blocking TAG synthesis restores autophagic flux. Remarkably, DAG supplementation also restores flux, increases PI abundance, and improves autophagosome completion despite elevated endogenous DAG levels. Together, these findings link Mdm1-dependent lipid homeostasis to autophagosome formation and suggest that coordination between membrane phospholipid synthesis and neutral lipid metabolism is important for efficient autophagy during starvation.

## Results

### Mdm1 is required for efficient starvation-induced autophagy

To determine whether Mdm1 is required for autophagy, we first measured bulk autophagic flux in wild-type (WT) and Δ*mdm1* cells using the green fluorescent protein (GFP)-Atg8 processing assay. GFP-Atg8 on the inner autophagosomal membrane is delivered to the vacuole with the autophagosome, where Atg8 is degraded and the more stable GFP moiety persists. The ratio of free GFP to total GFP (GFP-Atg8 plus free GFP) provides a measure of autophagic flux (Klionsky et al., 2021). We grew cells expressing endogenously tagged GFP-Atg8 to mid-logarithmic phase in synthetic complete medium (SCD) and shifted them to nitrogen-free medium (SD-N) for 3 h. Δ*mdm1* cells accumulated less free GFP than WT cells, and the ratio of free GFP to total GFP was significantly reduced (Fig. 1A and B).

**Figure 1.**
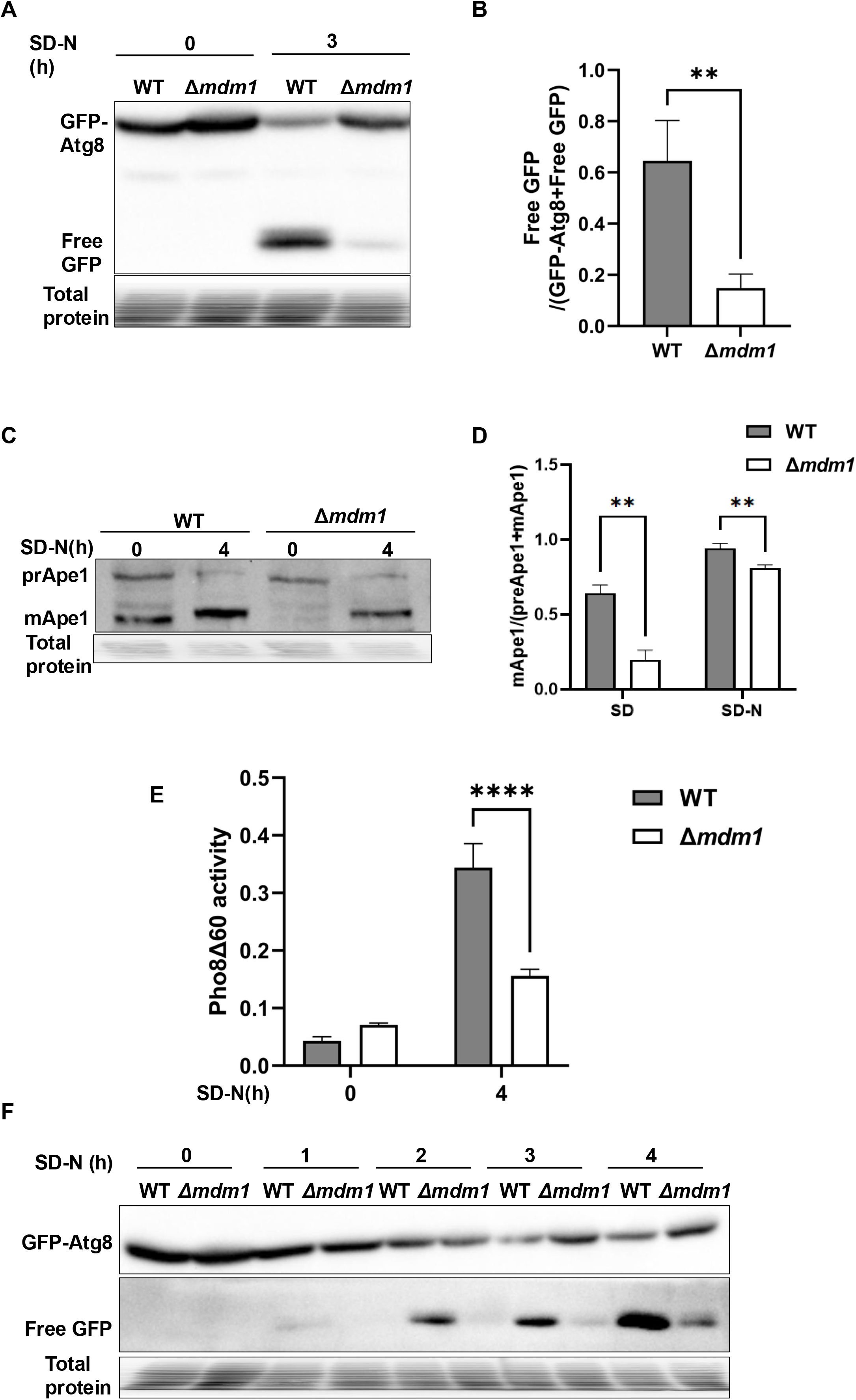
Mdm1 is required for efficient autophagic flux. **(A)** WT and Δ*mdm1* cells expressing endogenous GFP-Atg8 were grown to log phase in SCD and shifted to SD-N for 3 h. Whole-cell extracts were analysed by immunoblotting with anti-GFP antibodies. Total protein staining was used as a loading control. **(B)** Quantification of (A), expressed as free GFP/(GFP-Atg8 + free GFP). Data are mean ± SD of at least three independent experiments. Unpaired two-tailed Student’s t-test; **P < 0.01. **(C)** Ape1 maturation in WT and Δ*mdm1* cells before (0 h) and after 4 h of nitrogen starvation, detected by immunoblotting with anti-Ape1 antibodies. prApe1, precursor Ape1; mApe1, mature Ape1. Total protein staining was used as a loading control. **(D)** Quantification of (C), expressed as mApe1/(prApe1 + mApe1) in SD and SD-N. Data are mean ± SD of at least three independent experiments. Unpaired two-tailed Student’s t-test at each condition; **P < 0.01. **(E)** Pho8Δ60 activity in WT and Δ*mdm1* cells before (0 h) and after 4 h of nitrogen starvation. Data are mean ± SD of at least three independent experiments. Unpaired two-tailed Student’s t-test at 4 h; ****P < 0.0001. **(F)** GFP-Atg8 processing in WT and Δ*mdm1* cells at the indicated times of nitrogen starvation. Total protein staining was used as a loading control.

We next asked whether delivery of another autophagic cargo was also affected, by following maturation of the vacuolar hydrolase aminopeptidase I (Ape1). Ape1 is synthesized in the cytosol as a precursor, prApe1, and is processed to its mature form only after it reaches the vacuole. In growing cells, prApe1 is delivered by the selective cytoplasm-to-vacuole targeting (Cvt) pathway, whereas during nitrogen starvation it is carried mainly by autophagosomes (Klionsky et al., 2021). Because mature Ape1 (mApe1) migrates faster on SDS-PAGE, both forms can be measured on the same blot. After nitrogen starvation, Δ*mdm1* cells contained a smaller proportion of mApe1 than WT cells (Fig. 1C and D), showing that Mdm1 is also required for efficient delivery of autophagic cargo to the vacuole.

To measure flux by an independent route, we used the Pho8Δ60 assay, an enzymatic measure of bulk autophagy that does not rely on GFP-Atg8. Full-length Pho8 reaches the vacuole through the secretory pathway, directed by its N-terminal transmembrane domain. Pho8Δ60 lacks this domain and remains in the cytosol, so it reaches the vacuole, where it becomes active, only when it is engulfed by autophagosomes (Klionsky, 2007). Its activity is low in nutrient-replete cells and rises when autophagy is induced. After 4 h in SD-N, Pho8Δ60 activity was significantly lower in Δ*mdm1* cells than in WT cells (Fig. 1E). Together, these assays show that Mdm1 is required for efficient bulk autophagy during nitrogen starvation.

We wondered whether Δ*mdm1* cells were simply slower to start autophagy. To test this, we followed GFP-Atg8 processing at hourly intervals over 4 h of nitrogen starvation. Δ*mdm1* cells showed less processing than WT cells at every time point (Fig. 1F), indicating that flux is reduced throughout starvation rather than delayed.

We next asked whether the defect was specific to nitrogen starvation. Carbon starvation induces autophagy through a respiration-dependent pathway that is distinct from the response to nitrogen starvation (Adachi et al., 2017; Iwama & Ohsumi, 2019). Under this condition, Δ*mdm1* cells again processed less GFP-Atg8 than WT cells at both 3 h and 17 h (Fig. S1A and B). Rapamycin induces autophagy by inhibiting TORC1 directly, without removing any nutrient (Kamada et al., 2010). GFP-Atg8 processing was also reduced in rapamycin-treated Δ*mdm1* cells, although the reduction was smaller than under nitrogen starvation (Fig. S1C and D). These results show that Mdm1 is required for autophagy induced by different stimuli. Because rapamycin bypasses nutrient sensing, the rapamycin result also indicated that the defect in Δ*mdm1* cells lies downstream of nutrient sensing.

To test whether restoring Mdm1 corrects the defect, we expressed MDM1 from a plasmid in Δ*mdm1* cells. Plasmid-borne MDM1 did not restore GFP-Atg8 processing (Fig. S1E and F). A plasmid can fail to complement if Mdm1 above its normal level is itself inhibitory, so we asked whether an additional copy of MDM1 affects autophagy in WT cells, which already express endogenous Mdm1. Surprisingly, WT cells carrying the MDM1 plasmid showed significantly lower GFP-Atg8 processing than WT cells carrying the empty vector (Fig. S1G and H). An additional copy of MDM1 is therefore sufficient to reduce autophagic flux, indicating that autophagy is sensitive to MDM1 dosage. This sensitivity may explain why plasmid-borne MDM1 did not restore flux in Δ*mdm1* cells and indicates that efficient autophagic flux during starvation may require Mdm1 at its normal level.

### Mdm1 supports autophagosome formation

We next sought to identify the step of autophagy that requires Mdm1. A reduction in flux can arise in two ways. Autophagy may fail to be induced, or it may be induced normally while a later step, such as formation of the autophagosome or its fusion with the vacuole, is impaired. We began with induction. Nitrogen starvation and rapamycin inactivate TORC1, which leads to dephosphorylation of Atg13 and assembly of the Atg1 kinase complex that initiates phagophore formation (Kamada et al., 2010) (Fig. S2A). We expressed Atg13 from a multicopy plasmid under its own promoter and followed its phosphorylation by its mobility on immunoblots. After 30 min in SD-N, Atg13 was dephosphorylated to a similar extent in WT and Δ*mdm1* cells (Fig. S2B).

As a second measure of TORC1 activity, we followed the phosphorylation of Sch9, a direct TORC1 substrate (Urban et al., 2007). After chemical cleavage of Sch9-3×HA, the multiphosphorylated C-terminal fragment migrates more slowly than its dephosphorylated form, so the two can be distinguished on immunoblots. In SCD, WT and Δ*mdm1* cells contained similar proportions of multiphosphorylated Sch9 (Fig. S2C and D). After 30 min of nitrogen starvation, Sch9 shifted to its dephosphorylated form in both strains (Fig. S2C), and rapamycin produced the same shift (Fig. S2E). Cycloheximide, which increases TORC1 activity and Sch9 phosphorylation, served as a control for the hyperphosphorylated form (Fig. S2E). Total Sch9 was modestly lower in Δ*mdm1* cells, but the remaining protein responded to starvation as it did in WT cells. This experiment indicated that TORC1 signalling is intact in Δ*mdm1* cells and that the defect lies after induction.

With induction intact, we turned to the autophagic membranes themselves and asked whether they form and reach the vacuole in Δ*mdm1* cells. We imaged GFP-Atg8 at hourly intervals over 4 h of nitrogen starvation in cells treated with phenylmethylsulfonyl fluoride (PMSF). PMSF inhibits vacuolar serine proteases, so autophagic bodies that enter the vacuole are not degraded and their GFP-Atg8 signal builds up in the vacuolar lumen (Kawamata et al., 2022). GFP-Atg8 puncta outside the vacuole, in contrast, mark structures that have not yet fused. Over the time course, vacuolar GFP-Atg8 appeared in a growing proportion of WT cells, whereas significantly fewer Δ*mdm1* cells showed vacuolar signal (Fig. 2A and B). This was not because Δ*mdm1* cells lacked Atg8-positive membranes. More Δ*mdm1* cells than WT cells contained GFP-Atg8 puncta outside the vacuole, often close to the vacuolar membrane (Fig. 2C). Atg8-positive structures form in Δ*mdm1* cells, but they accumulate outside the vacuole. These structures could be autophagosomes that had not closed, or closed autophagosomes that had not fused with the vacuole.

**Figure 2.**
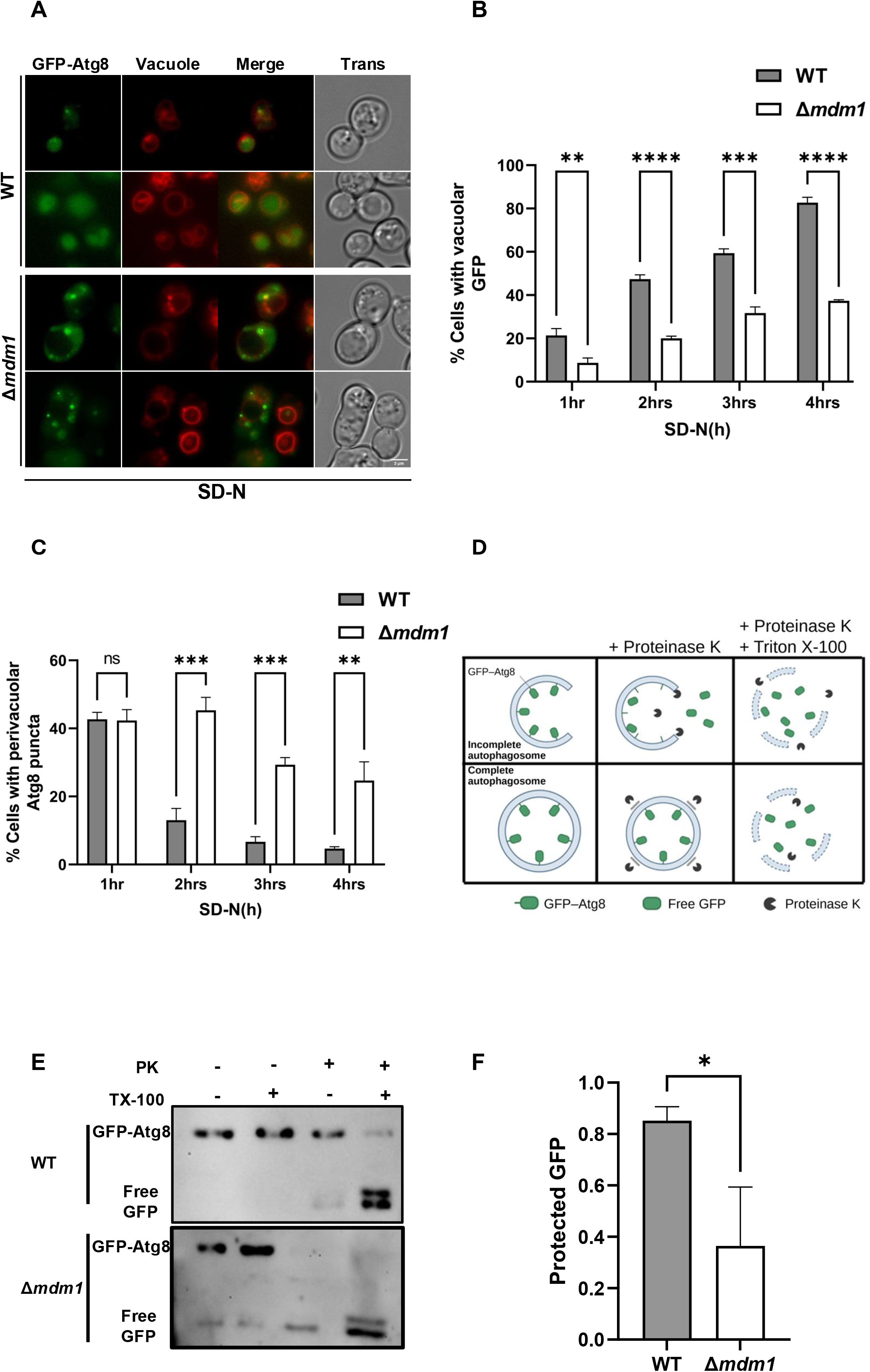
Mdm1 supports autophagosome formation. **(A)** WT and Δ*mdm1* cells expressing endogenous GFP-Atg8 were grown to log phase in SCD and shifted to SD-N supplemented with PMSF. Cells were imaged at hourly intervals during nitrogen starvation, and representative images are shown. Trans, transmitted light. Scale bar, 5 µm. **(B)** Percentage of WT and Δ*mdm1* cells with vacuolar GFP at the indicated times of nitrogen starvation. Data are mean ± SD of at least three independent experiments. Unpaired two-tailed Student’s t-test at each time point; **P < 0.01, ***P < 0.001, ****P < 0.0001. **(C)** Percentage of WT and Δ*mdm1* cells with perivacuolar GFP-Atg8 puncta at the indicated times of nitrogen starvation. Data are mean ± SD of at least three independent experiments. Unpaired two-tailed Student’s t-test at each time point; **P < 0.01, ***P < 0.001; ns, not significant. **(D)** Schematic of the protease-protection assay. Proteinase K degrades GFP-Atg8 that is exposed on incompletely closed structures, whereas GFP-Atg8 inside sealed autophagosomes is protected until the membranes are solubilized with Triton X-100. **(E)** WT and Δ*mdm1* cells in a Δ*pep4* background (labelled WT and Δ*mdm1*), expressing GFP-Atg8, were starved in SD-N for 3 h. Membrane fractions were treated with proteinase K (PK) with or without Triton X-100 (TX-100) and analysed by immunoblotting with anti-GFP antibodies. **(F)** Quantification of protease-protected GFP from (E) in WT and Δ*mdm1* cells in the Δ*pep4* background. Data are mean ± SD of at least three independent experiments. Unpaired two-tailed Student’s t-test; *P < 0.05.

To distinguish between these possibilities, we performed a protease-protection assay, which separates sealed autophagosomes from open phagophores (Nair et al., 2011) (Fig. 2D). We carried out the assay in a Δ*pep4* background, which lacks the major vacuolar protease activity. Membrane fractions from cells starved for 3 h were left untreated or were treated with proteinase K, Triton X-100, or both. GFP-Atg8 inside a sealed autophagosome is protected from proteinase K, whereas GFP-Atg8 on open structures is cleaved to free GFP. Detergent dissolves the membranes and exposes all of the GFP-Atg8. Δ*mdm1* cells contained significantly less protected GFP-Atg8 than WT cells (Fig. 2E and F), indicating that a larger fraction of their Atg8-positive structures had not sealed.

We next examined Atg8 lipidation as an independent measure of this step. During phagophore expansion, Atg8 is conjugated to phosphatidylethanolamine (PE), and as the autophagosome matures, Atg4 releases Atg8 from the outer membrane. Atg8-PE accumulates when autophagosome formation is impaired (Nair et al., 2012; Shpilka et al., 2015). After 4 h of nitrogen starvation, Δ*mdm1* cells contained more Atg8-PE relative to Atg8 than WT cells (Fig. S3A and B). Because lipidation depends on the upstream autophagy machinery, this result also shows that the early steps of autophagosome formation take place in Δ*mdm1* cells. We conclude that Mdm1 is required for autophagosome completion. In its absence, Atg8-positive membranes form and are lipidated, but a larger fraction of them fail to close and do not reach the vacuole.

Mdm1 resides at the NVJ, a contact site between the ER and the vacuole (Hariri et al., 2018, 2019), and the Atg8-positive structures that accumulated in Δ*mdm1* cells were often at the vacuolar periphery. We therefore asked whether Mdm1 lies near autophagic structures during starvation. Because endogenously tagged Mdm1 was too dim to image reliably under these conditions, we expressed Mdm1-mCherry from a plasmid in WT cells carrying endogenous GFP-Atg8. In nitrogen-starved cells, GFP-Atg8 puncta were frequently juxtaposed to Mdm1-mCherry at the vacuolar periphery, and line scans showed the two signals adjacent and in places partially overlapping (Fig. S3C). These images show that Mdm1 is positioned close to Atg8-positive structures during starvation.

### Mitophagy and lipophagy are reduced in Δ*mdm1* cells

Besides the NVJ, Mdm1 localizes to sites of LD biogenesis next to the vacuole (Hariri et al., 2018). Because Mdm1 sits at contacts between several organelles, we wondered whether its loss also affects the autophagic turnover of these organelles. Each reporter we used releases free GFP, or becomes enzymatically active, only after the organelle carrying it has been delivered to the vacuole. To examine mitophagy, we pregrew cells on lactate to increase their mitochondrial content and then starved them for nitrogen. After 6 h, processing of the mitochondrial matrix protein Idh1-GFP was reduced in Δ*mdm1* cells (Fig. S4A and B), and the activity of mitochondria-targeted Pho8Δ60 was also lower (Fig. S4C). Lipophagy, followed by processing of the LD protein Erg6-GFP after 5 h of starvation, was reduced as well (Fig. S4D and E). In contrast, processing of the ER membrane protein Sec63-GFP after 8 h of starvation did not differ between WT and Δ*mdm1* cells (Fig. S4F and G). Loss of Mdm1 reduces mitophagy and lipophagy, whereas ER-phagy reported by Sec63-GFP was unchanged under the conditions we tested. Because bulk autophagy is already impaired in Δ*mdm1* cells, the reduced mitophagy and lipophagy may follow from the general defect in autophagosome completion, and we focused the remaining experiments on bulk autophagy.

### Loss of Mdm1 alters phospholipid and neutral lipid homeostasis

The expansion and closure of the phagophore depend on a steady supply of newly synthesized phospholipid (Schütter et al., 2020a), and Mdm1 has been linked to LD biogenesis and lipid remodelling (Adebayo, Obaseki, Vasudeva, et al., 2026; Hariri et al., 2018, 2019; Obaseki et al., 2024; Paul et al., 2022). We asked whether lipid composition is altered in Δ*mdm1* cells. In yeast, PI, phosphatidylserine (PS) and PG are made from PA, whereas phosphatidylcholine (PC) and PE can also be made from DAG (Henry et al., 2012). Shotgun lipidomics of whole-cell extracts from nitrogen-starved cells showed that PI was significantly reduced in Δ*mdm1* cells, and PS was also significantly lower. PG tended to be lower, but the difference was not significant. PC, in contrast, was significantly increased, while PE was unchanged (Fig. 3A).

**Figure 3.**
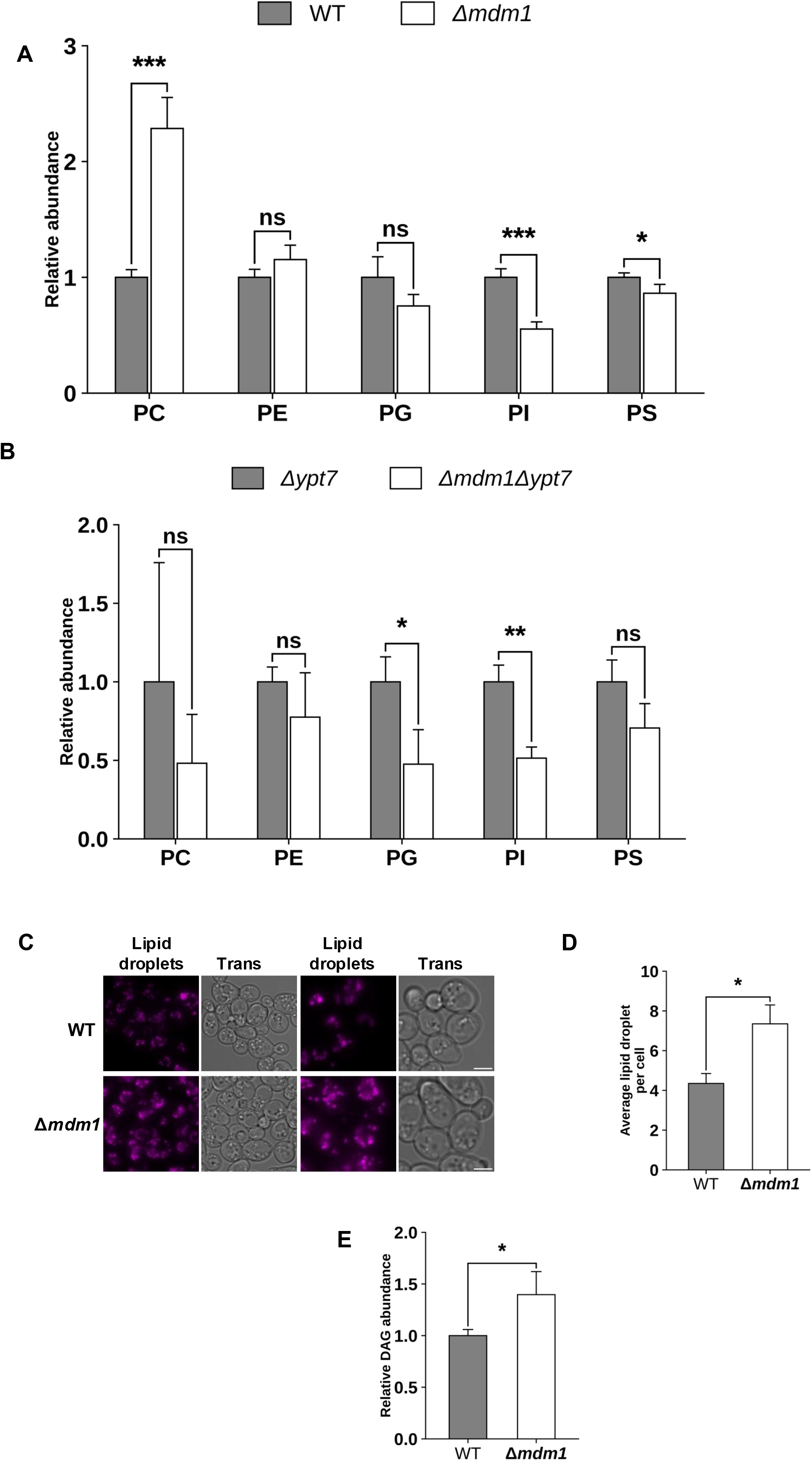
Loss of Mdm1 alters lipid homeostasis. (A) Relative abundance of phospholipid classes in whole-cell extracts of nitrogen-starved WT and Δ*mdm1* cells, determined by shotgun lipidomics and normalised to WT. Data are mean ± SD of four biological replicates. Unpaired two-tailed Student’s t-test for each lipid class; *P < 0.05, ***P < 0.001; ns, not significant. (B) Relative abundance of phospholipid classes in Atg8-enriched membranes isolated from nitrogen-starved Δ*ypt7* and Δ*mdm1* Δ*ypt7* cells, determined by shotgun lipidomics and normalised to Δ*ypt7*. Data are mean ± SD of three biological replicates. Unpaired two-tailed Student’s t-test for each lipid class; *P < 0.05, **P < 0.01; ns, not significant. (C) Lipid droplets in WT and Δ*mdm1* cells after 3 h of nitrogen starvation. Scale bar, 5 µm. (D) Average number of lipid droplets per cell from (C). Data are mean ± SD of at least three independent experiments. Unpaired two-tailed Student’s t-test; *P < 0.05. (E) Relative DAG abundance in whole-cell extracts of nitrogen-starved WT and Δ*mdm1* cells, determined by shotgun lipidomics and normalised to WT. Data are mean ± SD of four biological replicates. Unpaired two-tailed Student’s t-test; *P < 0.05.

Because whole-cell measurements combine all cellular membranes, we next examined the membranes of autophagosomes. In WT cells, autophagosomes fuse rapidly with the vacuole and are difficult to recover. We therefore used Δ*ypt7* cells, which lack the Rab GTPase Ypt7 and in which autophagosomes form but cannot fuse and accumulate in the cytoplasm. After 3 h of nitrogen starvation, we immunoisolated GFP-Atg8-positive membranes from Δ*ypt7* and Δ*ypt7* Δ*mdm1* cells. This procedure enriches for Atg8-positive structures and we refer to these preparations as Atg8-enriched membranes. Here again, PI was significantly reduced in Δ*ypt7* Δ*mdm1* cells, and PG was significantly reduced as well, whereas PC, PE and PS did not differ significantly (Fig. 3B). In this fraction, the PI acyl chains 16:1 and 18:0 were both reduced in Δ*ypt7* Δ*mdm1* cells (Fig. S5A). Important to note here that of all the phospholipids measured, PI was the only one reduced both in whole Δ*mdm1* cells and in Atg8-enriched membranes from Δ*ypt7* Δ*mdm1* cells.

Phospholipids and neutral lipids are made from shared intermediates, and loss of Mdm1 is known to increase cellular TAG (Hariri et al., 2018, 2019). We next asked whether neutral lipids accumulate in Δ*mdm1* cells during nitrogen starvation, when demand for membrane phospholipid is high. Δ*mdm1* cells stained with AutoDOT contained significantly more LDs per cell than WT cells after 3 h of nitrogen starvation (Fig. 3C and D). Lipidomic analysis showed that DAG was significantly increased, and several acyl chains of TAG were also significantly increased (Fig. 3E; Fig. S5B). Sterol esters (SE), measured by thin-layer chromatography after 4 h of starvation, were also significantly increased (Fig. S5C and D). During nitrogen starvation, Δ*mdm1* cells therefore accumulate DAG and storage lipids, while the PA-derived phospholipids PI and PS are depleted. We reasoned that this shortage of PA-derived phospholipids might contribute to their autophagy defect.

### Derepression of phospholipid synthesis restores PI and autophagic flux in Δ*mdm1* cells

If limited phospholipid synthesis contributes to the defect, increasing it should restore autophagy. To test this, we deleted *OPI1*. The transcriptional repressor Opi1 is held at the ER by binding PA. When PA levels fall, Opi1 moves into the nucleus and represses genes activated by Ino2-Ino4, including *INO1* and other phospholipid biosynthetic genes (Loewen et al., 2004). Deleting *OPI1* derepresses these genes. In Δ*opi1*Δ*mdm1* cells, PI was significantly increased and returned to WT levels (Fig. 4A). Importantly, these cells also processed GFP-Atg8 at WT levels after 4 h in SD-N (Fig. 4B and C). OPI1 deletion also significantly increased PG in Δ*mdm1* cells, while PC and PE were unchanged (Fig. 4A). In the WT background, OPI1 deletion increased each phospholipid class measured, as expected from derepression of phospholipid biosynthetic genes (Henry et al., 2012) (Fig. 4A). Derepressing phospholipid synthesis therefore restored both PI and autophagic flux in Δ*mdm1* cells. Because PI has well-established roles in autophagosome biogenesis (Baba & Balla, 2020; Laczkó-Dobos et al., 2021; Schütter et al., 2020a), we focused on PI.

**Figure 4.**
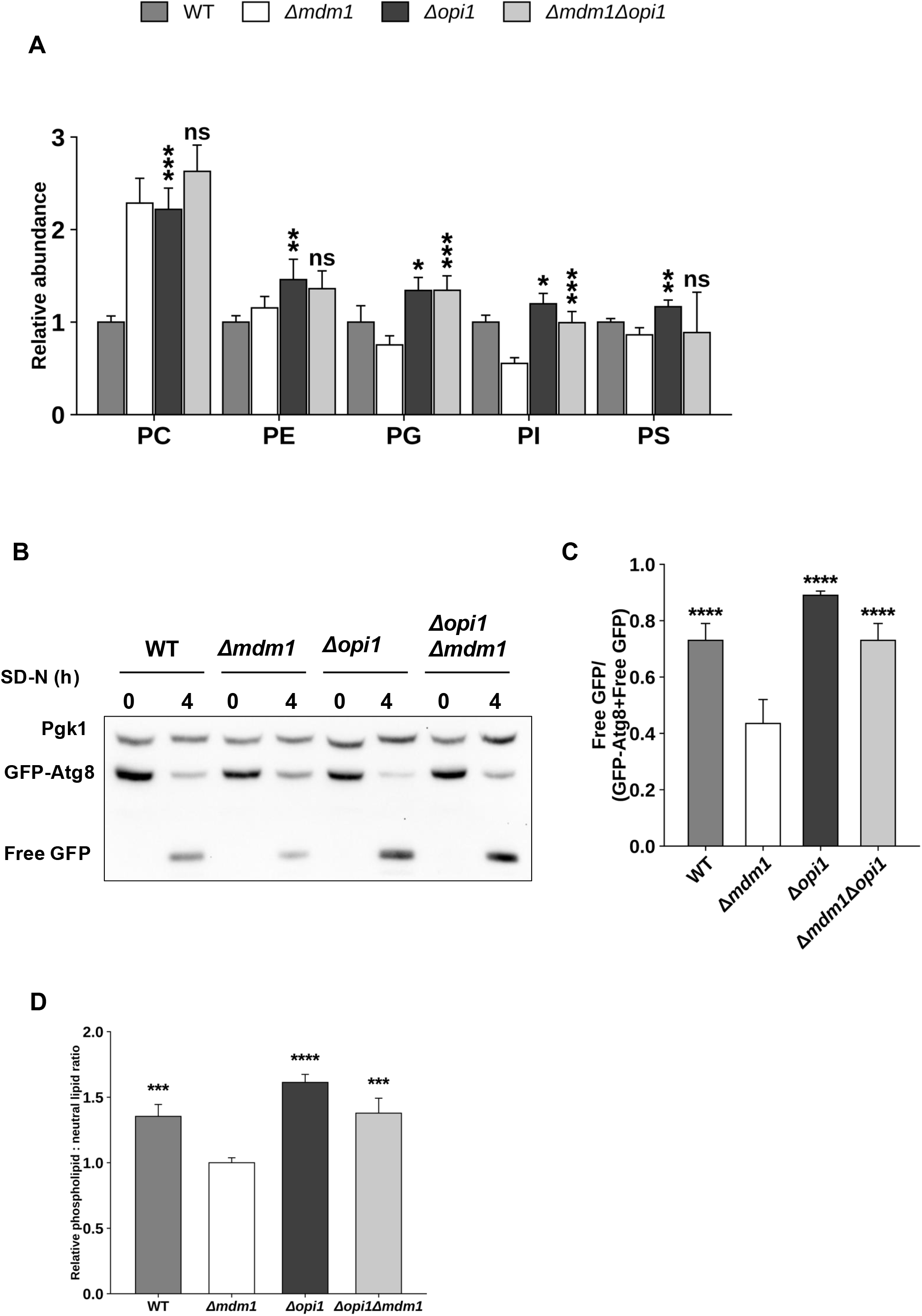
*OPI1* deletion rebalances phospholipid homeostasis in Δ*mdm1* cells and restores autophagic flux. (A) Relative abundance of phospholipid classes in whole-cell extracts of nitrogen-starved WT, Δ*mdm1*, Δ*opi1* and Δ*mdm1* Δ*opi1* cells, determined by shotgun lipidomics and normalised to WT. Asterisks above Δ*opi1* show comparison with WT, and asterisks above Δ*mdm1* Δ*opi1* show comparison with Δ*mdm1*. Data are mean ± SD of four biological replicates. Unpaired two-tailed Student’s t-test for each lipid class; *P < 0.05, **P < 0.01, ***P < 0.001; ns, not significant. (B) GFP-Atg8 processing in WT, Δ*mdm1*, Δ*opi1* and Δ*opi1* Δ*mdm1* cells before (0 h) and after 4 h of nitrogen starvation, analysed by immunoblotting with anti-GFP antibodies. Pgk1 was used as a loading control. (C) Quantification of (B), expressed as free GFP/(GFP-Atg8 + free GFP). Asterisks show comparison with Δ*mdm1*. Data are mean ± SD of at least three independent experiments. Unpaired two-tailed Student’s t-test; ****P < 0.0001. (D) Ratio of total phospholipid to neutral lipid in WT, Δ*mdm1*, Δ*opi1* and Δ*opi1* Δ*mdm1* cells, normalised to Δ*mdm1*. Asterisks show comparison with Δ*mdm1*. Data are mean ± SD of four biological replicates. Unpaired two-tailed Student’s t-test; ***P < 0.001, ****P < 0.0001.

PI is made from inositol, so we next asked whether a shortage of inositol caused the low PI in Δ*mdm1* cells. If it did, adding inositol should restore flux, as *OPI1* deletion did. However, supplementation with 75 µM myo-inositol did not restore GFP-Atg8 processing in Δ*mdm1* cells (Fig. S6A and B). PC and PI are the most abundant phospholipids of yeast autophagosomal membranes, followed by PE (Schütter et al., 2020b), so we also supplied inositol, choline and ethanolamine together (ICE) to support synthesis of all three. ICE did not restore flux either (Fig. S6C and D).

We also asked whether Δ*mdm1* cells have a defect in inositol or PI synthesis. Myo-inositol-3-phosphate synthase (MIPS), the *INO1* product that catalyzes the first step of inositol synthesis, was significantly more abundant in Δ*mdm1* cells than in WT cells (Fig. S6E and F). MIPS increased strongly when *OPI1* was deleted and became nearly undetectable after ICE supplementation (Fig. S6G), showing that *INO1* still responds normally to Opi1 and to inositol in Δ*mdm1* cells. Expression of *PIS1*, which encodes PI synthase, was also unchanged (Fig. S6H). Together, these results indicate that low PI in Δ*mdm1* cells is not caused by reduced inositol synthesis or by reduced *PIS1* expression.

If phospholipid homeostasis limits autophagy in Δ*mdm1* cells, impairing it further should make the defect worse. To test this, we deleted *OPI3*, which encodes the methyltransferase that converts phosphatidylmonomethylethanolamine, produced from PE by Cho2, to PC. Opi3 is required to maintain phospholipid balance during autophagosome completion (Polyansky et al., 2022). As predicted, Δ*opi3* Δ*mdm1* cells had less autophagy than either single mutant (Fig. S5E and F). This result indicates that autophagy in Δ*mdm1* cells remains sensitive to further perturbation of phospholipid synthesis.

We also compared the ratio of total phospholipid to neutral lipid. This ratio was significantly lower in Δ*mdm1* cells than in WT cells, and *OPI1* deletion returned it to WT levels (Fig. 4D). Because the recovery of autophagic flux in Δ*opi1* Δ*mdm1* cells coincided with a WT-level balance between membrane and storage lipids, we next manipulated neutral lipid synthesis directly

### TAG and DAG metabolism modulate autophagic flux in Δ*mdm1* cells

Phospholipids and neutral lipids branch from two shared intermediates, PA and DAG (Fig. S8A). Pah1 dephosphorylates PA to produce DAG, which can then be acylated to TAG or phosphorylated back to PA by the DAG kinase Dgk1 (Carman & Han, 2006; Han et al., 2008). In yeast, almost all TAG is made by two enzymes, Dga1, which uses acyl-CoA as the acyl donor, and Lro1, which transfers an acyl chain from a phospholipid (Oelkers et al., 2000, 2002; Sandager et al., 2002). We reasoned that if TAG synthesis in Δ*mdm1* cells consumes intermediates that would otherwise support membrane synthesis, blocking it should improve autophagy. Interestingly, deletion of *DGA1* and *LRO1* restored GFP-Atg8 processing in Δ*mdm1* cells to WT levels after 4 h of nitrogen starvation (Fig. 5A and B), showing that blocking TAG synthesis restores autophagy in Δ*mdm1* cells.

**Figure 5.**
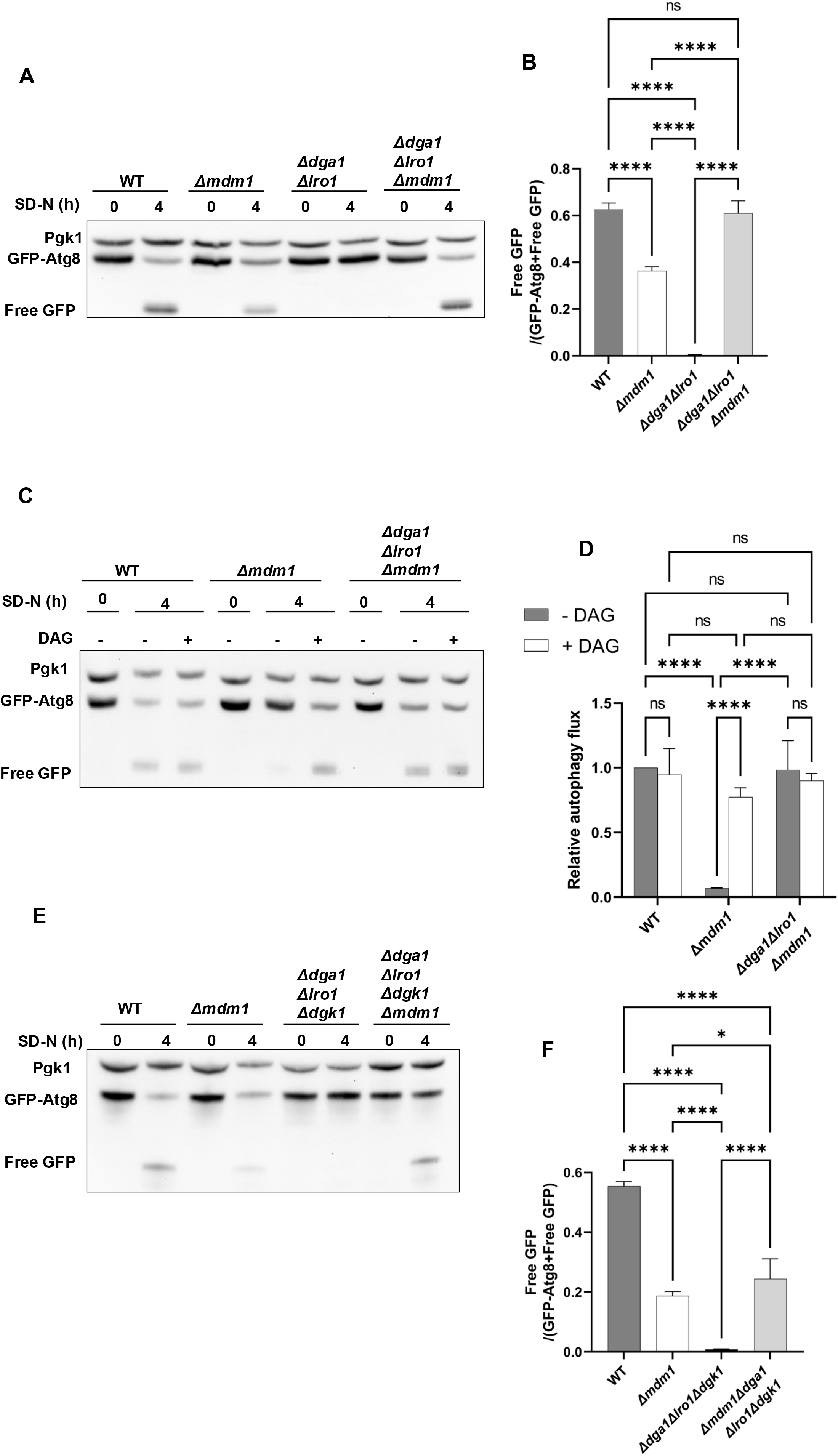
Blocking TAG synthesis or supplying DAG restores autophagic flux in Δ*mdm1* cells. **(A)** GFP-Atg8 processing in WT, Δ*mdm1*, Δ*dga1* Δ*lro1* and Δ*dga1* Δ*lro1* Δ*mdm1* cells before (0 h) and after 4 h of nitrogen starvation, analysed by immunoblotting with anti-GFP antibodies. Pgk1 was used as a loading control. **(B)** Quantification of (A), expressed as free GFP/(GFP-Atg8 + free GFP). Data are mean ± SD of at least three independent experiments. One-way ANOVA with Tukey’s multiple comparisons test; ****P < 0.0001; ns, not significant. **(C)** GFP-Atg8 processing in WT, Δ*mdm1* and Δ*dga1* Δ*lro1* Δ*mdm1* cells before (0 h) and after 4 h of nitrogen starvation, in the absence (−) or presence (+) of DAG. Pgk1 was used as a loading control. **(D)** Quantification of (C), expressed as autophagic flux relative to WT without DAG, which was set to 1. Data are mean ± SD of at least three independent experiments. One-way ANOVA with Tukey’s multiple comparisons test; ****P < 0.0001; ns, not significant. **(E)** GFP-Atg8 processing in WT, Δ*mdm1*, Δ*dga1* Δ*lro1* Δ*dgk1* and Δ*mdm1* Δ*dga1* Δ*lro1* Δ*dgk1* cells before (0 h) and after 4 h of nitrogen starvation. Pgk1 was used as a loading control. **(F)** Quantification of (E), expressed as free GFP/(GFP-Atg8 + free GFP). Data are mean ± SD of at least three independent experiments. One-way ANOVA with Tukey’s multiple comparisons test; *P < 0.05, ****P < 0.0001.

To determine whether this effect was specific to TAG, we blocked synthesis of the other storage lipid, SE, by deleting *ARE1* and *ARE2*, which encode the sterol acyltransferases. Rather than improving autophagy, loss of SE synthesis further reduced GFP-Atg8 processing in Δ*mdm1* cells (Fig. S5G and H). This result indicates that the rescue is specific to blocking TAG synthesis.

Blocking TAG synthesis spares both of its substrates, fatty acids and DAG, and either could account for the rescue. We next wanted to delineate which of them is important for autophagy in Δ*mdm1* cells. We first tested fatty acids. Cells transferred to SD-N containing 0.1 mM of a mixture of the free fatty acids (FFA) palmitic, stearic and myristic acid showed increased AutoDOT staining within 1 h, confirming that the fatty acids were taken up and stored (Fig. S7A). FFA did not, however, restore GFP-Atg8 processing in Δ*mdm1* cells (Fig. S7B and C).

We next asked whether DAG could restore autophagy in Δ*mdm1* cells, adding it across a range of concentrations (0.005 to 1.45 mM) at the onset of nitrogen starvation. Low concentrations of DAG increased GFP-Atg8 processing, with the strongest effect at 0.025 mM, whereas concentrations of 0.75 mM and above reduced processing below the untreated level (Fig. S7D), indicating that only low-level supplementation of DAG supports autophagy in Δ*mdm1* cells. We therefore used 0.025 mM DAG in the experiments that followed. At this concentration, DAG restored GFP-Atg8 processing in Δ*mdm1* cells to WT levels, in contrast to FFA supplementation (Fig. 5C and D). Strikingly, this rescue occurred even though total DAG is already elevated in Δ*mdm1* cells (Fig. 3E). Also, DAG supplementation did not further increase GFP-Atg8 processing in WT cells or in Δ*dga1* Δ*lro1* Δ*mdm1* cells, in which TAG synthesis is already blocked (Fig. 5C and D). Blocking TAG synthesis and supplying DAG therefore both restore autophagy in Δ*mdm1* cells, and their effects are not additive, consistent with both acting by increasing the DAG available for autophagy in these cells.

DAG spared by the TAG block could support autophagy in two ways. It could be converted by Dgk1 back to PA, from which PI, PS and PG are made, or it could act downstream of this conversion. To distinguish between these, we deleted *DGK1* in Δ*dga1* Δ*lro1* Δ*mdm1* cells. Loss of Dgk1 reduced the rescue but did not abolish it. GFP-Atg8 processing in Δ*dga1* Δ*lro1* Δ*dgk1* Δ*mdm1* (4KO) cells was higher than in Δ*mdm1* cells but remained below the WT level (Fig. 5E and F), and Pho8Δ60 activity in the same strains gave the same result (Fig. S7E). Together, these results indicate that conversion of DAG to PA by Dgk1 accounts for part of the rescue, but not all of it.

If DAG availability limits autophagy in Δ*mdm1* cells, reducing DAG production should make the defect worse. Pah1 produces DAG from PA, and deletion of *PAH1* further reduced GFP-Atg8 processing in Δ*mdm1* cells, to a level below that of either Δ*mdm1* or Δ*pah1* cells (Fig. S8B and C). Together, these experiments indicate that autophagy in Δ*mdm1* cells is limited not by the total amount of DAG but by the DAG that remains available for use, in part through its conversion back to PA by Dgk1. We next asked how DAG supplementation changes the lipid composition of Δ*mdm1* cells, and whether it restores autophagosome completion.

### DAG supplementation restores autophagosome completion in Δ*mdm1* cells

To determine how exogenous DAG changes the lipid profile of Δ*mdm1* cells, we starved WT and Δ*mdm1* cells with or without DAG and analysed them by shotgun lipidomics, as in Fig. 3A. In Δ*mdm1* cells, DAG significantly increased PI and PE and significantly decreased PS, while PC and PG were unchanged (Fig. 6A). The PI acyl chains that increased with DAG were the same ones reduced in Atg8-enriched membranes from Δ*ypt7* Δ*mdm1* cells (Fig. S8D; Fig. S5A). The added DAG did not accumulate in Δ*mdm1* cells. Total DAG and TAG were unchanged in these cells, whereas both increased significantly in DAG-treated WT cells (Fig. S7I), indicating that Δ*mdm1* cells use the added DAG differently from WT cells. As a result, the ratio of phospholipid to neutral lipid increased significantly in DAG-supplemented Δ*mdm1* cells, as it did after *OPI1* deletion (Fig. S8E), although it did not reach the WT level. DAG also increased this ratio in WT cells (Fig. S8F). In Δ*mdm1* cells, DAG addition therefore raised PI and PE and shifted the balance from storage lipids toward membrane phospholipids.

**Figure 6.**
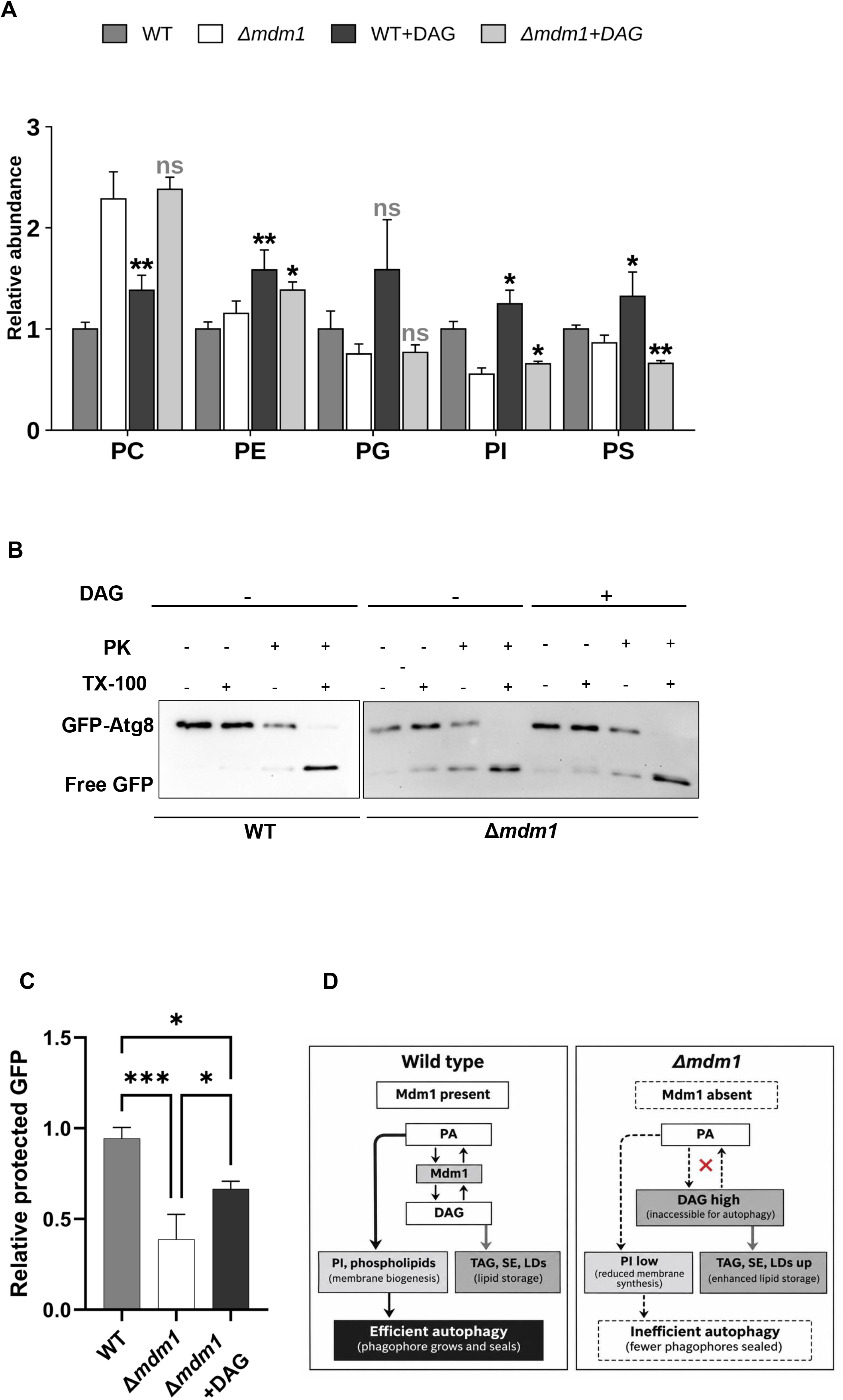
DAG supplementation promotes autophagosome completion in Δ*mdm1* cells. **(A)** Relative abundance of phospholipid classes in whole-cell extracts of nitrogen-starved WT and Δ*mdm1* cells with or without DAG, determined by shotgun lipidomics and normalised to WT. Asterisks above WT+DAG show comparison with WT. Asterisks above Δ*mdm1*+DAG show comparison with Δ*mdm1*. Data are mean ± SD of four biological replicates. Unpaired two-tailed Student’s t-test for each lipid class; *P < 0.05, **P < 0.01; ns, not significant. **(B)** WT and Δ*mdm1* cells in a Δ*pep4* background (labelled WT and Δ*mdm1*), expressing GFP-Atg8, were starved in SD-N for 3 h in the absence (−) or presence (+) of DAG. Membrane fractions were treated with proteinase K (PK) with or without Triton X-100 (TX-100) and analysed by immunoblotting with anti-GFP antibodies. **(C)** Quantification of protease-protected GFP from (B), relative to WT. Data are mean ± SD of at least three independent experiments. One-way ANOVA with Tukey’s multiple comparisons test; *P < 0.05, ***P < 0.001. **(D)** Working model. In WT cells, Mdm1 favours the use of DAG for membrane phospholipid synthesis, which supports efficient autophagy. In Δ*mdm1* cells, DAG is diverted to storage lipids, PI is reduced and fewer phagophores are sealed. Arrow thickness indicates the proposed relative flow of lipid.

Finally, we asked whether DAG corrects the step at which our results show that autophagy fails in Δ*mdm1* cells, autophagosome completion. WT and Δ*mdm1* cells were starved for 3 h with or without DAG and analysed by protease protection. Interestingly, DAG supplementation significantly increased the level of cargo protection of autophagosomes of Δ*mdm1* cells (Fig. 6B and C). Together, these data support a model of lipid allocation in which Mdm1 helps direct DAG-derived precursors toward the membrane lipid synthesis needed to complete autophagosomes during nitrogen starvation (Fig. 6D).

## Discussion

Efficient execution of autophagy, particularly the formation and maturation of the double-membrane autophagosome, requires precise coordination between membrane supply, lipid metabolism, and signaling pathways that operate under conditions of nutrient stress (Reggiori & Klionsky, 2013). While much attention has focused on the core autophagy machinery that initiates phagophore formation, how cells prioritize lipid resources to support autophagosome completion remains incompletely understood. In this study, we identify Mdm1 as a critical regulator of autophagosome maturation that acts by coordinating lipid availability during starvation, revealing an unappreciated layer of metabolic control of autophagy.

Our results show that Δ*mdm1* cells have a defect in starvation-induced autophagy and that this defect originates from inefficient autophagosome formation. Induction was intact in these cells, as TORC1 was inactivated and Atg13 was dephosphorylated. Atg8-positive structures formed and Atg8 was lipidated, but fewer of these structures were sealed and reached the vacuole. Flux was restored by three manipulations of lipid metabolism: derepression of phospholipid synthesis, a block in TAG synthesis and supplementation of DAG. These rescues indicate that the autophagy defect of Δ*mdm1* cells depends on lipid metabolism.

How might Mdm1 be coordinating this lipid supply required for autophagosome formation? Previous studies have reported that Mdm1 localizes at ER-vacuole-lipid droplet contact sites, where it regulates lipid-droplet biogenesis during nutrient stress (Hariri et al., 2018, 2019). In our study, exogenously expressed Mdm1 localized adjacent to Atg8-positive structures during starvation, suggesting that Mdm1 may regulate lipid availability locally at sites of autophagosome formation. Its PXA and PXC domains are predicted to facilitate lipid transfer, raising a possibility that Mdm1 directs lipid intermediates away from TAG storage and toward phospholipid synthesis needed for phagophore expansion and closure (Aboumourad & Hariri, 2026; Paul et al., 2022). This model is conceptually similar to Atg2, which transfers lipids to growing autophagosomal membranes and also contributes to lipid-droplet homeostasis (Elhan et al., 2025; Valverde et al., 2019). Mdm1 may therefore function as a contact-site lipid regulator that helps allocate lipid intermediates according to metabolic demand during starvation.

A model role of Mdm1 in local lipid allocation may also explain why both MDM1 deletion and overexpression impair autophagy. Mdm1 overexpression and loss elevates lipid droplet levels (Hariri et al., 2018). A similar dosage dependence has been reported for ATG2A, where DGAT2 relocalization to lipid droplets depends on ATG2A abundance and activity (Elhan et al., 2025). These findings suggest that lipid transfer proteins must be maintained within an appropriate level to efficiently coordinate lipid metabolism. Accordingly, either insufficient or excess Mdm1 may perturb local lipid partitioning and limit lipid precursors available for autophagosome formation.

The reduction in PI is particularly notable because PI and its phosphorylated derivatives contribute to multiple stages of autophagy, including recruitment of autophagy factors, membrane growth identity, membrane trafficking, and autophagosome formation(Baba & Balla, 2020; Kurokawa et al., 2019). Interesting for us, was that the PI species restored by DAG supplementation overlapped with species reduced in autophagosome-enriched fractions from Mdm1-deficient cells.

DAG emerges as a central metabolic branchpoint in this model because it can be committed to TAG synthesis or converted by Dgk1 into PA, which feeds phospholipid production. One limitation our study did not resolve was why supplementing DAG, or inhibiting TAG synthesis to genetically spare it, rescues autophagy in Δ*mdm1* cells that already accumulates DAG. This paradox may be reconciled if cellular DAG is not a single pool, but it is partitioned into spatially and distinct metabolic populations, and this has been reported in literature (Ganesan et al., 2015). Loss of mdm1 may be biasing DAG towards storage committed pools and consumed for TAG synthesis, at the expense of membrane synthesis pool.

DAG may also influence autophagy independently of its metabolic conversion, as it can alter membrane curvature and act as a lipid-signaling molecule in other systems (Eichmann & Lass, 2015). The ability of both genetic inhibition of TAG synthesis and exogenous DAG supplementation to rescue autophagy in Δ*mdm1* cells suggests that DAG availability is limiting when Mdm1 is absent. Although the partial requirement for Dgk1 supports a contribution from DAG conversion to PA, to probably support PI synthesis.

A further open question is why loss of Mdm1 preferentially impairs mitophagy and lipophagy while leaving ER-phagy intact. Mdm1 has previously been reported to be dispensable for packaging of the ER into autophagosomes, a step that instead depends on the lipid transporter, Vps13 (Chen et al., 2020). An explanation could be that mitophagy and lipophagy depend on different lipid metabolic rewiring. Mitophagy has been shown to be unusually sensitive to phospholipid metabolism. Mss4, which makes PI 4,5-bisphosphate, is required for mitophagy but not for bulk autophagy (K. Wang et al., 2012), and loss of Opi3 or of the Nem1-Spo7 phosphatase impairs mitophagy more strongly than bulk autophagy (Sakakibara et al., 2015; Xu & Okamoto, 2018). A reduced PI pool could limit the phosphoinositides that mitophagy depends on. Studies have also shown that lipophagy rely on membrane microdomains enriched in raft forming lipids (C. W. Wang et al., 2014), of which the proteomes of NVJ have been shown to play a role in (Sakuragi et al., 2023). These possibilities will need to be addressed by additional experiments.

Finally, our data extend the emerging role of SNX-RGS proteins in autophagy. The Drosophila SNX-RGS protein Snazarus and its human orthologue SNX25 modulate autophagic flux, while loss of mammalian SNX13 and SNX14 are associated with abnormal autophagic-vacuole accumulation (Akizu et al., 2015; Lauzier et al., 2022; Zheng et al., 2006). Our work identifies Mdm1 as a yeast SNX-RGS protein that promotes starvation-induced autophagy by coordinating phospholipid and neutral-lipid metabolism at membrane contact sites. Important questions remain, whether Mdm1 directly transfers specific lipid species to forming autophagosomes and which lipid species. Addressing this question will define how Mdm1 preserves the local lipid balance required for autophagosome completion during nutrient stress.

## Materials and methods

### Yeast Strains, media and Culture Conditions

All yeast strains used are listed in Table S1 and plasmids, in Table S2. Gene deletions were generated following standard yeast knock-in/knock-out protocols (Gietz & Schiestl, 2007). Strains were selected using antibiotics or dropout selection media. Yeast cells were cultured at 30°C in either yeast peptone dextrose (YPD: 1% yeast extract, 2% peptone, 2% glucose) or synthetic complete medium (SCD: 0.67% yeast nitrogen base, 2% glucose, supplemented with complete amino acids and vitamins), as indicated. For mitochondrial proliferation, cells were grown in lactate-containing medium (YPL: 1% yeast extract, 2% peptone, 2% lactic acid) for 18 h. To induce autophagy, mid log phase cells were washed twice with sterile water and transferred into nitrogen starvation medium with glucose (SD-N: 0.17% yeast nitrogen base without ammonium sulfate and amino acids, 2% glucose) or treated with rapamycin (200ng/mL) for the indicated durations.

### Pho8Δ60 assay

Pho8Δ60 activity measurement were performed as previously described (Gómez Sánchez et al., 2018). 5 OD_600_ equivalents of cells were lysed in 400 μL ice-cold lysis buffer (20 mM Pipes, pH 6.8, 0.5% Triton X-100, 50 mM KCl, 100 mM potassium acetate, 10 mM MgCl_2_, 10 μM ZnSO_4_, and 2 mM PMSF) by vortexing with glass beads for 3-5 min at 4°C. Lysates were centrifuged at 13,000 × g for 5 min at 4°C. Subsequently, 100 μL of the resulting supernatant was mixed with 400 μL alkaline phosphatase reaction buffer (250 mM Tris-HCl, pH 8.5, 0.4% Triton X-100, 10 mM MgCl_2_, 10 μM ZnSO_4_, and 1.25 mM p-nitrophenyl phosphate) prewarmed to 37°C. Samples were incubated at 37°C for 20 min, after which 500 μL of 1 M glycine (pH 11.0) was added to stop the reaction. Samples were briefly centrifuged at 13,000 × g for 2 min, and the absorbance of the supernatant was measured at 400 nm.

### Protease-Protection Assay

Cells (60-100 OD600 units) were harvested by centrifugation at 3000 revolutions/min for 5 min. Cells were washed once with H2O, resuspended in 25 mL freshly prepared DTT buffer (10 mM Tris, pH 9.4, 10 mM DTT), and incubated at 30°C for 10 min in a water bath. Cells were then collected by centrifugation, washed twice with 5 mL H2O, and resuspended in 5 mL SP buffer (1 M sorbitol, 20 mM PIPES, pH 6.8). Zymolyase 20T (0.74 mg per OD unit, pre-dissolved in SP buffer) was added to the suspension, and cells were incubated at 30°C for 45 min with gentle agitation every 15 min to form spheroplasts. The spheroplasts were carefully layered onto 5 mL of 1.8 M sorbitol in a 15-mL tube, centrifuged at 2000 revolutions/min for 10 min, and the supernatant was discarded. The pelleted spheroplasts were gently resuspended in 5 mL PS200 buffer (20 mM PIPES, pH 6.8, 200 mM sorbitol, 5 mM MgCl_2_). Spheroplasts were then homogenized with 10 strokes in a pre-chilled Dounce homogenizer. Homogenates were transferred to pre-cooled 15-mL tubes and centrifuged at 400 × g for 10 min at 4°C twice to remove unbroken cells and cellular debris. The clarified supernatant was divided into four 1-mL aliquots and centrifuged at 5000 × g for 5 min at 4°C. Each pellet was resuspended in 300 µL PS200 buffer containing one of the following conditions: (1) PS200 alone, (2) 0.2% Triton X-100, (3) 40 µg/mL proteinase K, or (4) 0.2% Triton X-100 plus 40 µg/mL proteinase K, and incubated on ice for 30 min. To terminate the reaction, 33 µL of trichloroacetic acid (TCA) was added. Samples were mixed thoroughly by vortexing and incubated on ice for 30 min. After centrifugation at 13,000 revolutions/min for 10 min, pellets were washed twice with acetone, air-dried, resuspended in 50 µL MURB buffer (50 mM sodium phosphate buffer, pH 7.0, 25 mM MES, 1% SDS [w/v], 3 M urea, 1 mM NaN3, 1% β-mercaptoethanol, 0.01% bromophenol blue), and heated at 55°C for 15 min.

### Immunoprecipitation of GFP-Atg8 membranes

Cells expressing GFP-Atg8 were cultured in SCD medium and shifted to SD-N medium for 3 h. Harvested yeast pellets were resuspended in four pellet volumes of homogenization buffer (10 mM Tris-HCl, pH 7.4, 0.25 M sucrose) supplemented with 2 mM PMSF and protease inhibitor cocktail. Cells were homogenized by bead-beating (glass beads, 4 cycles of 45 s ON and 1 min OFF, at 4°C). Unbroken cells and nuclei were removed by centrifugation at 12,000 × g for 5 min at 4°C. Equal amounts of the cleared homogenate were ultracentrifuged (speed; 90,000 rpm, 30 min, 4°C) to obtain membrane fractions. Pellets were resuspended in homogenization buffer and incubated with an anti-GFP antibody (Sigma, 11814460001) on ice for 1 h, followed by overnight incubation with Protein G agarose beads (Sigma, 11719416001) with rotation at 4°C. The beads were washed three times with washing buffer (10 mM Tris-HCl, pH 7.5, 150 mM NaCl, 0.5 mM EDTA). Immunoprecipitation was validated by eluting proteins and performing western blot analysis. Subsequently, beads from parallel samples (without elution) were submitted for lipidomics analysis.

### qPCR

Cells were harvested by centrifugation (4000□ ×□ g, 5□min), supernatant aspirated and stored at −80□°C. After thawing on ice, total RNA was isolated using the Nucleospin RNA Plus kit (Takara Bio USA, 740984) according to the manufacturer’s protocol. Briefly, cell pellets were resuspended in 350□µL LBP buffer (Nucleospin RNA Plus kit) and ∼0.5 volume 0.5mm acid washed glass beads before lysis using bead beater (4□×□1□min cycles at maximum speed, with 2 min on ice between cycles). RNA was further purified using the same kit per the supplier’s instructions. cDNA synthesis was carried out with the PrimeScript RT reagent kit with genomic DNA eraser (Takara Bio USA, #RR092A). qPCR was performed in 10□µL reactions using PowerUp SYBR Green Master Mix (Thermo Fisher, #A25777); transcript levels were normalized to the housekeeping gene taf10. Primer efficiencies were validated empirically.

### Quantitative image analysis

Live-cell imaging of yeast cultures was performed using an EVOS M5000 microscope (Thermo Fisher Scientific) at room temperature. Yeast cells were grown to the desired optical density under the specified growth conditions. Images were analyzed using Fiji (ImageJ). Lipid droplets were quantified by counting per cell using the Multi-Point Tool. Identical acquisition and processing parameters were applied across samples.

### LD staining

For LD staining, cells were grown as described above, pelleted by centrifugation (4000 x g, 5 minutes), supernatant aspirated, and incubated for 5 min with monodansylpentane (MDH; Abcepta, #SM1000b). After incubation, cells were washed once with water, pelleted again, and imaged. For vacuolar staining, cells were incubated for at least 30 min with the styryl dye FM 4-64 (Invitrogen, T13320) at a final concentration of 1 μM directly in the culture medium before collection for microscopic analysis.

### Lipid extraction, quantitative lipidomics of yeast cells and isolated autophagosome

All the solvents used were HPLC or LC/MS grade and purchased from Sigma-Aldrich (St Louis, MO, USA). Splash® Lipidomix® standards were purchased from Avanti (Alabaster, AL, USA). The lipid liquid-liquid extractions (LLE) were performed in 16×100mm glass tubes with PTFE-lined caps (Fisher Scientific, Pittsburg, PA, USA. Aliquots corresponding to 250K cells were transferred to fresh glass tubes for LLE. Lipids were extracted using a modified methyl tert-butyl ether (mTBE) method (Matyash et al., 2008). Briefly, 1mL of water, 1 mL of methanol, and 2mL of mTBE were added to the glass tubes containing the sample. The mixture was vortexed and centrifuged at 2,671 g for 5 min. The organic phase was transferred to a fresh glass tube and spiked with 20µL of a 1:5 diluted Splash® Lipidomix® standards mixture (Avanti Polar Lipids, Alabaster, AL). Samples were dried under nitrogen and resuspended in 400 µl of hexane.

The lipid profile analysis was performed by LC-MS/MS using a SCIEX QTRAP 6500+ (SCIEX, Framingham, MA) coupled to a Shimadzu LC-30AD (Shimadzu, Columbia, MD) high-performance liquid chromatography (HPLC) system. Separation was achieved in a 150×2.1 mm, 5 µm Supelco Ascentis silica column (Supelco, Bellefonte, PA. Samples were injected at a flow rate of 0.3 ml/min, starting with 2.5% solvent B (mTBE) and 97.5% Solvent A (hexane). The gradient was as follows: Solvent B was increased to 5% over 3 min and then to 60% over 6 min. Solvent B was decreased to 0% during 30 sec while Solvent C (90:10 (v/v) isopropanol-water) was set at 20% and increased to 40% during the following 11 min. Solvent C is increased to 44% over 6 min and then to 60% over 50 sec. The system was held at 60% solvent C for 1 min prior to re-equilibration at 2.5% of solvent B for 5 min at a 1.2 mL/min flow rate. Solvent D [95:5 (v/v) acetonitrile-water with 10 mM Ammonium acetate] was infused post-column at 0.03 ml/min. The column oven temperature was maintained at 25°C. Data was acquired in both positive and negative ionization mode using multiple reaction monitoring (MRM). The LC-MS/MS data was analyzed using MultiQuant software (SCIEX). The identified lipid species were normalized to their corresponding internal standard.

### Thin layer chromatography

#### Lipid extraction

Lipids were extracted using a modified Folch method optimized for neutral lipids. Cell pellets were weighed, resuspended in sterile, deionized water, and disrupted with acid-washed glass beads using a mini bead beater (4 × 2 min, maximum speed, 4°C). Chloroform:methanol (2:1, v/v) was added, vortexed (5 minutes at room temperature), and transferred to a glass tube (Group A). The original tube was rinsed with methanol before combining with Group A. Additional chloroform was added, vortexed (1 minute), and centrifuged (2000 × g, 5 minutes, room temperature). The resulting lower phase was transferred to a new tube (Group B). Group A was re-extracted with 2:1 chloroform:methanol and the lower phase was combined with Group B. Sterile, deionized water was added to assist in phase separation as needed. Group B was washed twice with 1 M potassium chloride, vortexed, and centrifuged (2000 × g, 5 min, room temperature), discarding upper phases. The final organic phase was collected, dried under argon gas, and resuspended in chloroform (1 mg wet weight/µL). Samples (4–6 µL) were used for TLC analysis.

Silica gel plates were prebaked (145°C, 30 min), then marked with 18 lanes (1.5 cm outer margins, 1 cm between lanes) using a pencil. The development chamber was equilibrated with the appropriate solvent system for neutral lipids: hexane: diethyl ether: acetic acid (80:20:10). Samples were applied and plates were developed at room temperature until lipid bands were fully separated. After air drying for 20 min, plates were sprayed with 3% copper (II) acetate in 8% phosphoric acid and charred at 145 °C for 1-8 hours to visualize bands; spraying and heating were repeated as needed.

#### TLC quantification

Stained TLC plates were scanned and then processed for quantification using Fiji (ImageJ). Each plate contained a serial dilution of a standardized neutral lipid mixture. The standard mixture was prepared in chloroform to a final concentration of 10 mg/mL. The standardized neutral lipid mixture was used to create a standard curve in which the x axis displayed the calculated lipid mass in micrograms, and the y-axis displayed the estimated band intensity. Statistical analysis was performed in GraphPad Prism.

#### Western blot analysis

For GFP-Atg8 cleavage assays, yeast whole-cell extracts were prepared using a modified alkaline lysis method based on (Kushnirov, 2000). Cell pellets were first resuspended in 100□μl of distilled water, followed by addition of 100□μl of 0.2□M sodium hydroxide (NaOH). After briefly vortexing, samples were incubated at room temperature for 5 minutes. Cells were then centrifuged at 18,000□×□g for 30 seconds, and the supernatant was aspirated.

The resulting pellets were resuspended in 1X Laemmli buffer, with the volume adjusted according to the culture’s optical density unit to normalize total protein input. Samples were boiled at 95□°C for 5 minutes before centrifugation at 18,000□×□g for 1 minute.

For SDS-PAGE analysis, 6-10□μl of cleared lysate was loaded per lane in 10% polyacrylamide gels and run at 115V for 40-60 minutes. Proteins were semi-dry transferred to a PVDF membrane before blocking in 5% non-fat milk in TBST (20□mM Tris-HCl pH 7.5, 150□mM NaCl, 0.1% Tween-20). Membranes were probed with αGFP primary antibody diluted in blocking solution (1:1000; Millipore-Sigma #11814460001). Detection was performed with HRP-conjugated secondary antibodies and ECL (BioRad Clarity™ Western ECL, #1705061) reagents, and signal was imaged on an iBright FL1500 system (Thermo Fisher). Total protein staining (No-Stain™ Protein Labelling Reagent, Fisher Scientific # A44449) or anti-Pgk1, served as the loading control. Atg13 phosphorylation was as described in Miller-Fleming et al., 2014 and Sch9 phosphorylation was as described in Murley et al., 2015.

## Author contributions

E. Obaseki and H. Hariri, conception and design of the study, data interpretation, drafting and revising the manuscript. E. Obaseki, D. Adebayo, M. Rahal, imaging, image quantification, immunoblot analysis, data analysis, figure preparation, critical reading of the manuscript, generating yeast strains, technical and editorial assistance. G. Vale and J. G. McDonald, phospholipid and neutral lipid analysis.

## Acknowledgements

We thank Dr. Daniel Klionsky for sharing strains, plasmids and antibodies (anti-Ape1, anti-Atg13), UTSW Medical Center Lipid and Metabolite Mass Spectrometry Facility for lipidomics expertise, and members of the Miriam Greenberg laboratory for sharing supplies. H. Hariri is supported by the National Institutes of Health National Institute of General Medical Sciences (R35GM150892), Wayne State University start-up funds and a University Research Award, a Karmanos Cancer Institute Strategic Research Initiative Grant (KCI-SRIG), and the Richard Barber Interdisciplinary Research Program. The Microscopy, Imaging and Cytometry Resources Core is supported in part by an NIH Karmanos Cancer Institute Center Grant (P30 CA22453) and Wayne State University.

The authors declare no competing financial interest.

---

**Figure S1. Δ*mdm1* cells have reduced autophagic flux under carbon starvation and rapamycin treatment, and autophagy is sensitive to *MDM1* levels.**

**(A)** WT and Δ*mdm1* cells expressing endogenous GFP-Atg8 were grown to log phase in SCD and shifted to carbon-starvation medium for 0, 3 or 17 h. Whole-cell extracts were analysed by immunoblotting with anti-GFP antibodies. Total protein staining was used as a loading control.

**(B)** Quantification of (A), expressed as free GFP/(GFP-Atg8 + free GFP). Data are mean ± SD of two biological replicates.

**(C)** WT and Δ*mdm1* cells expressing endogenous GFP-Atg8 were grown to log phase in SCD and treated with 200 nM rapamycin for 3 h. Pgk1 was used as a loading control.

**(D)** Quantification of (C), expressed as free GFP/(GFP-Atg8 + free GFP). Data are mean ± SD of at least three independent experiments. Unpaired two-tailed Student’s t-test; **P < 0.01.

**(E)** WT cells carrying an empty vector (EV), and Δ*mdm1* cells carrying EV or a plasmid expressing *MDM1* (p*MDM1*), all expressing GFP-Atg8, were grown to log phase in SD-URA and shifted to SD-N for 4 h. Pgk1 was used as a loading control.

**(F)** Quantification of (E), expressed as free GFP/(GFP-Atg8 + free GFP). Data are mean ± SD of at least three independent experiments. One-way ANOVA with Tukey’s multiple comparisons test; ***P < 0.001, ****P < 0.0001; ns, not significant.

**(G)** WT cells carrying EV or p*MDM1*, expressing GFP-Atg8, were grown to log phase in SD-URA and shifted to SD-N for 4 h. Total protein staining was used as a loading control.

**(H)** Quantification of (G), expressed as free GFP/(GFP-Atg8 + free GFP). Data are mean ± SD of at least three independent experiments. Unpaired two-tailed Student’s t-test; *P < 0.05.

**Figure S2. TORC1 signalling and autophagy induction are intact in Δ*mdm1* cells.**

**(A)** Schematic of TORC1 signalling during nitrogen starvation or rapamycin treatment. Active TORC1 phosphorylates Atg13, which prevents assembly of the Atg1–Atg13–Atg17 complex, and phosphorylates Sch9, which promotes translation and ribosome biogenesis. When TORC1 is inhibited, Atg13 is dephosphorylated and autophagy is initiated.

**(B)** Atg13 phosphorylation in WT and Δ*mdm1* cells expressing Atg13 from a multicopy plasmid under its endogenous promoter. Cells were grown to log phase in SCD and shifted to SD-N for 30 min, and whole-cell extracts were analysed by immunoblotting with anti-Atg13 antibodies. P-Atg13, phosphorylated Atg13.

**(C)** Phosphorylation of Sch9 in WT and Δ*mdm1* cells expressing Sch9-3×HA, grown in SCD and shifted to SD-N for 30 min. The Sch9 C-terminal fragment was released by chemical cleavage and analysed by immunoblotting with anti-HA antibodies. P-Sch9, multiphosphorylated Sch9, which migrates more slowly. Total protein staining was used as a loading control.

**(D)** Ratio of P-Sch9 to total Sch9 in SCD, quantified from (C). Data are mean ± SD of at least three independent experiments. Unpaired two-tailed Student’s t-test; ns, not significant.

**(E)** Sch9 phosphorylation in WT and Δ*mdm1* cells grown in SCD, treated with rapamycin (Rap), or treated with cycloheximide (CHX), analysed as in (C).

**Figure S3. Δ*mdm1* cells accumulate lipidated Atg8, and Mdm1 is proximal to Atg8-positive structures during nitrogen starvation.**

**(A)** Atg8 lipidation in WT and Δ*mdm1* cells after 4 h of nitrogen starvation, analysed by immunoblotting with anti-Atg8 antibodies. Atg8-PE, phosphatidylethanolamine-conjugated Atg8.

**(B)** Ratio of Atg8-PE to Atg8 from (A), normalised to WT. Data are mean ± SD of at least three independent experiments. Unpaired two-tailed Student’s t-test; *P < 0.05.

**(C)** WT cells co-expressing endogenous GFP-Atg8 and plasmid-borne Mdm1-mCherry (Mdm1-mCH) were grown to log phase in SCD, shifted to SD-N and imaged during nitrogen starvation. Five representative cells are shown. Trans, transmitted light. Scale bar, 5 µm. For each cell, a line scan of fluorescence intensity along the indicated line is shown on the right (GFP-Atg8 in green, Mdm1-mCH in red).

**Figure S4. Loss of Mdm1 reduces mitophagy and lipophagy but not ER-phagy.**

**(A)** WT and Δ*mdm1* cells expressing endogenous Idh1-GFP were grown to log phase in SCD and shifted to SD-N for 6 h. Whole-cell extracts were analysed by immunoblotting with anti-GFP antibodies. Total protein staining was used as a loading control.

**(B)** Quantification of (A), expressed as free GFP/(Idh1-GFP + free GFP). Data are mean ± SD of at least three independent experiments. Unpaired two-tailed Student’s t-test; ****P < 0.0001.

**(C)** Mito-Pho8Δ60 activity in WT and Δ*mdm1* cells before (0 h) and after 6 h of nitrogen starvation. Data are mean ± SD of at least three independent experiments. Unpaired two-tailed Student’s t-test at 6 h; *P < 0.05.

**(D)** WT and Δ*mdm1* cells expressing endogenous Erg6-GFP were grown to log phase in SCD and shifted to SD-N for 5 h, and analysed as in (A).

**(E)** Quantification of (D), expressed as free GFP/(Erg6-GFP + free GFP) relative to WT. Data are mean ± SD of at least three independent experiments. Unpaired two-tailed Student’s t-test; *P < 0.05.

**(F)** WT and Δ*mdm1* cells expressing endogenous Sec63-GFP were grown to log phase in SCD and shifted to SD-N for 8 h, and analysed as in (A).

**(G)** Quantification of (F), expressed as free GFP/(Sec63-GFP + free GFP), with WT set to 1. Data are mean ± SD of at least three independent experiments. Unpaired two-tailed Student’s t-test; ns, not significant.

**Figure S5. Loss of Mdm1 alters PI, TAG and sterol esters, and deleting *OPI3* or *ARE1/ARE2* worsens autophagy in** Δ***mdm1* cells.**

**(A)** PI species in Atg8-enriched membranes isolated from nitrogen-starved Δ*ypt7* and Δ*mdm1*Δ*ypt7* cells, analysed by shotgun lipidomics. Values are relative to Δ*ypt7*, with 16:1 and 18:0 PI shown. Data are mean ± SD of three independent preparations (n = 3). Unpaired two-tailed Student’s t-test for each species; **P < 0.01.

**(B)** TAG acyl chains (14:0, 16:0, 16:1, 18:0 and 18:1) in whole-cell lipid extracts from WT and Δ*mdm1* cells, analysed by shotgun lipidomics. Values are relative to WT. Data are mean ± SD of four independent cultures (n = 4). Unpaired two-tailed Student’s t-test for each acyl chain; *P < 0.05, ***P < 0.001; ns, not significant.

**(C)** Sterol esters in WT and Δ*mdm1* cells after 4 h in SD-N, separated by thin-layer chromatography. Each lane is an independent culture.

**(D)** Quantification of (C). Data are mean ± SD of three independent cultures. Unpaired two-tailed Student’s t-test; *P < 0.05.

**(E)** WT, Δ*mdm1*, Δ*mdm1*Δ*opi3* and Δ*opi3* cells expressing GFP-Atg8 were grown to log phase in SCD (0 h) and shifted to SD-N for 4 h. Whole-cell extracts were immunoblotted with anti-GFP and anti-Pgk1 antibodies. Pgk1 was the loading control.

**(F)** Quantification of (E), expressed as free GFP/(GFP-Atg8 + free GFP) at 4 h. Data are mean ± SD of at least three independent experiments. One-way ANOVA with Tukey’s multiple comparisons test; *P < 0.05, **P < 0.01, ***P < 0.001; ns, not significant.

**(G)** WT, Δ*mdm1*, Δ*are1*Δ*are2* and Δ*are1*Δ*are2*Δ*mdm1* cells expressing GFP-Atg8 were grown and analysed as in (E).

**(H)** Quantification of (G), expressed as in (F). Data are mean ± SD of at least three independent experiments. One-way ANOVA with Tukey’s multiple comparisons test; ***P < 0.001, ****P < 0.0001; ns, not significant.

**Figure S6. Inositol or ICE supplementation does not restore autophagy in Δ*mdm1* cells, and MIPS is increased in Δ*mdm1* cells.**

**(A)** WT and Δ*mdm1* cells expressing GFP-Atg8 were grown to log phase in SCD (0 h) and shifted to SD-N for 4 h. Where indicated, the medium contained 75 µM myo-inositol. Whole-cell extracts were immunoblotted with anti-GFP and anti-Pgk1 antibodies. Pgk1 was the loading control.

**(B)** Quantification of (A) at 4 h, expressed as free GFP/(GFP-Atg8 + free GFP). Data are mean ± SD of at least three independent experiments. Each strain was compared with Δ*mdm1* using an unpaired two-tailed Student’s t-test; **P < 0.01; ns, not significant.

**(C)** WT and Δ*mdm1* cells expressing GFP-Atg8 were grown with or without ICE (75 µM inositol, 1 mM choline and 50 mM ethanolamine), then analysed as in (A).

**(D)** Quantification of (C), as in (B). Data are mean ± SD of at least three independent experiments. Each strain was compared with Δ*mdm1* using an unpaired two-tailed Student’s t-test; **P < 0.01; ns, not significant.

**(E)** MIPS (Ino1) in WT and Δ*mdm1* cells, detected by immunoblotting with anti-MIPS antibodies. Total protein staining was the loading control. Each lane is an independent culture.

**(F)** Quantification of (E), expressed as MIPS/total protein stain (TPS). Data are mean ± SD of three independent cultures. Unpaired two-tailed Student’s t-test; ****P < 0.0001.

**(G)** MIPS in WT, Δ*mdm1*, Δ*opi1* and Δ*mdm1*Δ*opi1* cells, and in WT and Δ*mdm1* cells grown with ICE, detected as in (E). The numbers below each lane are MIPS/total protein from this blot.

(H) *PIS1* mRNA in WT and Δ*mdm1* cells, measured by RT-qPCR and expressed relative to WT. Data are mean ± SD of at least three independent experiments. Unpaired two-tailed Student’s t-test; ns, not significant.

**Figure S7. Effects of free fatty acids, DAG and loss of DAG consumption on autophagy in**Δ***mdm1* cells.**

**(A)** Δ*mdm1* cells were grown for 1 h with free fatty acids (FFA; 0.1 mM each of palmitic, stearic and myristic acid in 1% Tween-40) or with vehicle (ethanol). Lipid droplets were stained with MDH. Two representative fields are shown per condition. Trans, transmitted light. Scale bar, 3 µm.

**(B)** WT and Δ*mdm1* cells expressing GFP-Atg8 were grown to log phase in SCD (0 h) and shifted to SD-N for 4 h, with or without FFA as in (A). Whole-cell extracts were immunoblotted with anti-GFP and anti-Pgk1 antibodies, and Pgk1 was used as the loading control.

**(C)** Quantification of (B) at 4 h, expressed as free GFP/(GFP-Atg8 + free GFP). Data are mean ± SD of at least three independent experiments. One-way ANOVA with Tukey’s multiple comparisons test; ***P < 0.001; ns, not significant.

**(D)** GFP-Atg8 processing in WT cells, and in Δ*mdm1* cells supplemented with increasing concentrations of DAG (0–1.45 mM), during 4 h in SD-N, analysed as in (B). Values below the blot are free GFP/(GFP-Atg8 + free GFP) relative to WT at 4 h, from this blot.

**(E)** Pho8Δ60 activity in WT, Δ*mdm1*, Δ*dga1*Δ*lro1*Δ*dgk1* and Δ*mdm1*Δ*dga1*Δ*lro1*Δ*dgk1* cells in SCD and after 4 h in SD-N. Data are mean ± SD of at least three independent experiments. SD-N values were compared by one-way ANOVA with Tukey’s multiple comparisons test; ****P < 0.0001.

**Figure S8. Effects of *PAH1* deletion and DAG supplementation on autophagy and lipid balance in Δ*mdm1* cells.**

**(A)** Schematic of phospholipid and neutral lipid synthesis in yeast.

**(B)** WT, Δ*mdm1*, Δ*pah1* and Δ*mdm1*Δ*pah1* cells expressing GFP-Atg8 were grown to log phase in SCD (0 h) and shifted to SD-N for 4 h. Whole-cell extracts were immunoblotted with anti-GFP and anti-Pgk1 antibodies, and Pgk1 was used as the loading control.

**(C)** Quantification of (B) at 4 h, expressed as free GFP/(GFP-Atg8 + free GFP). Data are mean ± SD of at least three independent experiments. One-way ANOVA with Tukey’s multiple comparisons test; **P < 0.01, ****P < 0.0001.

**(D)** 16:1, 18:0 and 18:1 PI species in whole-cell lipid extracts from WT, Δ*mdm1* and Δ*mdm1* + DAG cells, measured by shotgun lipidomics. Values are relative to Δ*mdm1*. Data are mean ± SD of four independent cultures (n = 4). Each strain was compared with Δ*mdm1* by unpaired two-tailed Student’s t-test for each species; *P < 0.05, ***P < 0.001, ****P < 0.0001; ns, not significant.

**(E)** TAG and DAG in whole-cell lipid extracts from WT, Δ*mdm1*, WT + DAG and Δ*mdm1* + DAG cells, measured by shotgun lipidomics. Values are relative to WT. Data are mean ± SD of four independent cultures (n = 4). WT + DAG was compared with WT, and Δ*mdm1* + DAG with Δ*mdm1*, by unpaired two-tailed Student’s t-test; **P < 0.01; ns, not significant.

**(F)** Phospholipid:neutral lipid ratio of the cells in (D), relative to Δ*mdm1*. Data are mean ± SD of four independent cultures (n = 4). Unpaired two-tailed Student’s t-test against Δ*mdm1*; ***P < 0.001, ****P < 0.0001.

