## Supplemental figures for "ER-vacuole tether Mdm1 coordinates lipid homeostasis to support autophagosome formation"

### Fig. S1

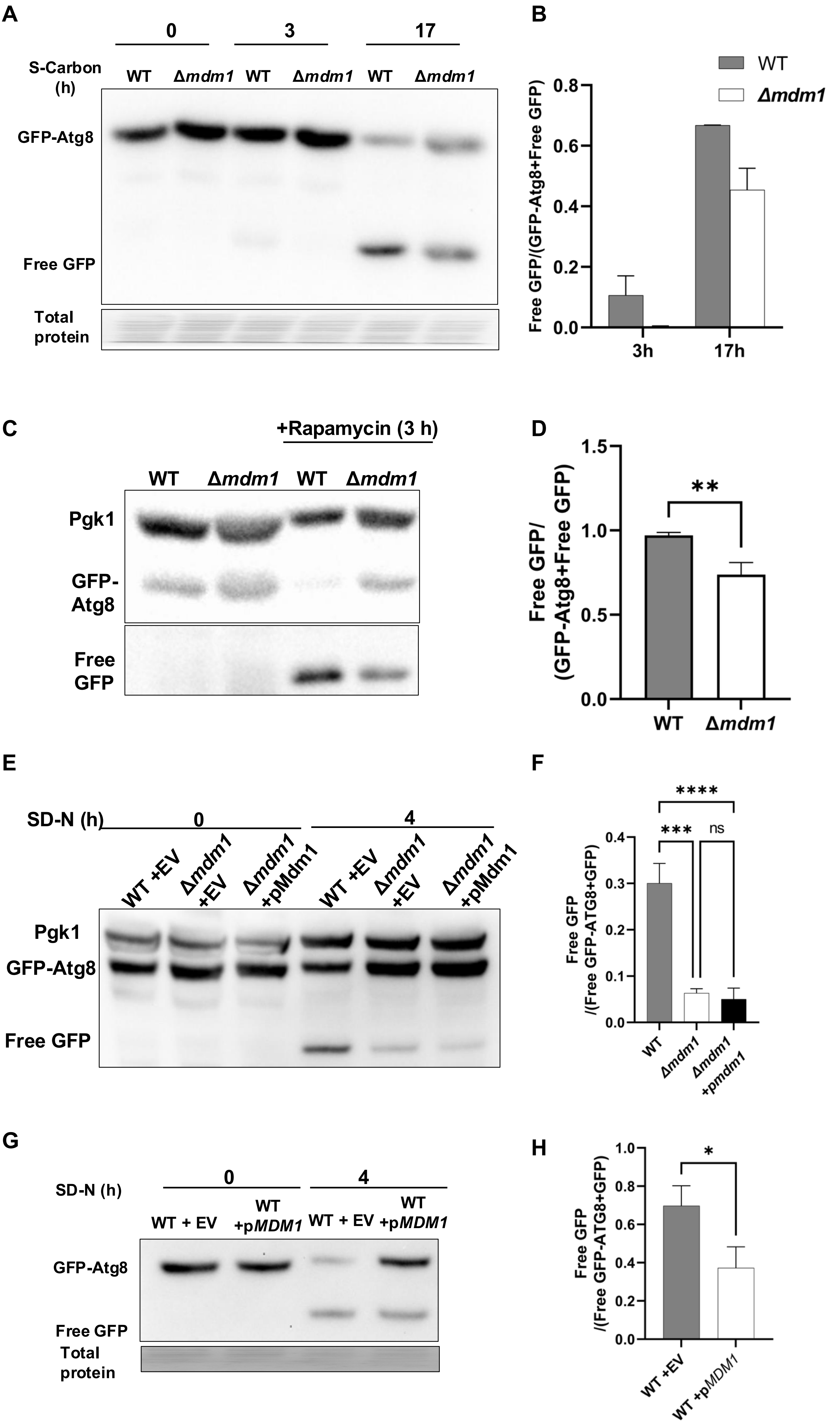
A
S-Carbon (h)
0
3
17
WT
Δmdm1
WT
Δmdm1
WT
Δmdm1
GFP-Atg8
Free GFP
Total protein
B
WT
Δmdm1
Free GFP/(GFP-Atg8+Free GFP)
0.0
0.2
0.4
0.6
0.8
3h
17h
C
+Rapamycin (3 h)
WT
Δmdm1
WT
Δmdm1
Pgk1
GFP-Atg8
Free GFP
D
Free GFP/(GFP-Atg8+Free GFP)
0.0
0.5
1.0
1.5
**
WT
Δmdm1
E
SD-N (h)
0
4
WT +EV
Δmdm1 +EV
Δmdm1 +pMdm1
WT +EV
Δmdm1 +EV
Δmdm1 +pMdm1
Pgk1
GFP-Atg8
Free GFP
F
Free GFP /(Free GFP-ATG8+GFP)
0.0
0.1
0.2
0.3
0.4
****
***
ns
WT
Δmdm1
Δmdm1 +pmdm1
G
SD-N (h)
0
4
WT + EV
WT +pMDM1
WT + EV
WT +pMDM1
GFP-Atg8
Free GFP
Total protein
H
Free GFP /(Free GFP-ATG8+GFP)
0.0
0.2
0.4
0.6
0.8
1.0
*
WT +EV
WT +pMDM1

### Fig. S2

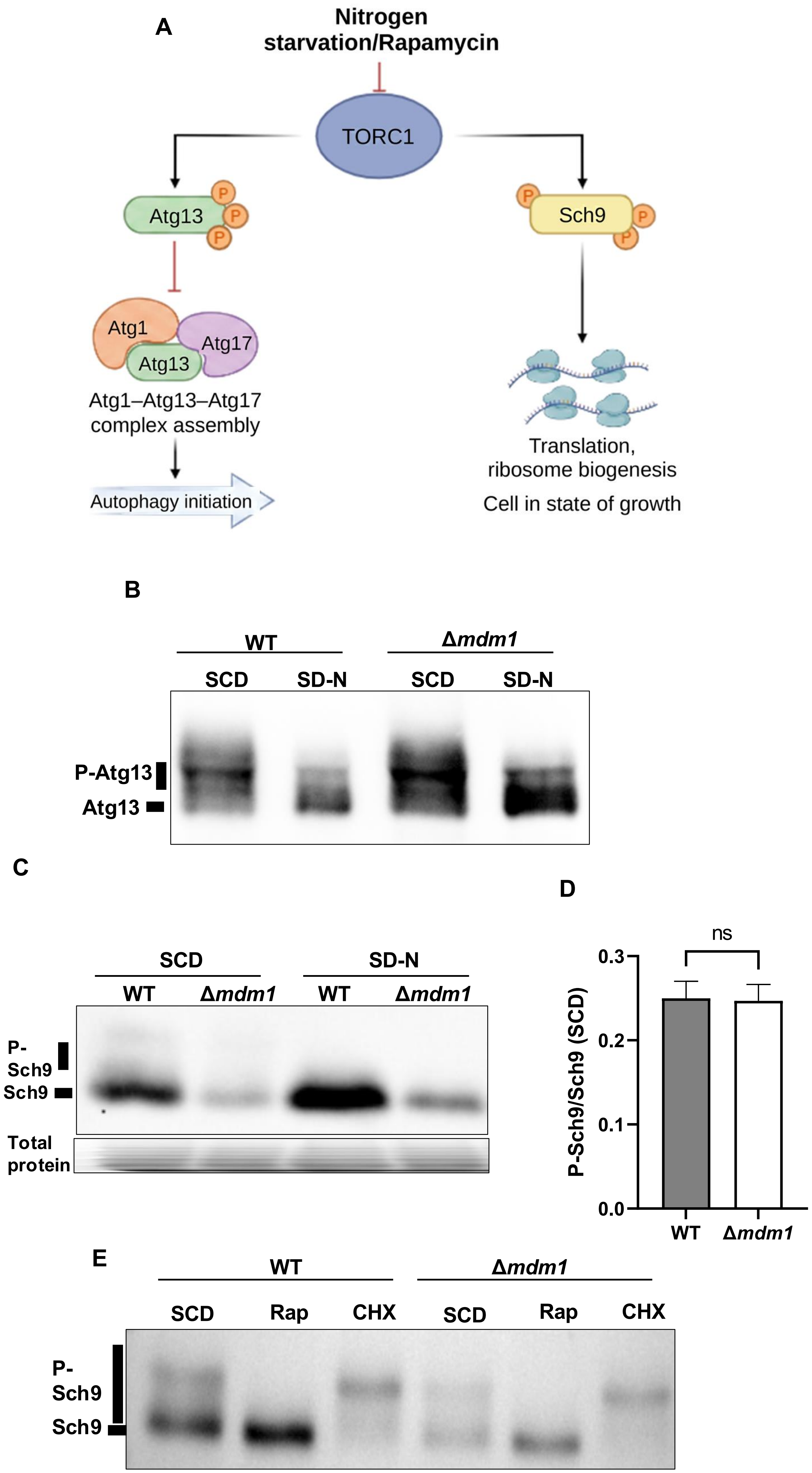

A
Nitrogen starvation/Rapamycin
TORC1
Atg13
P
Sch9
Atg1
Atg17
Atg13
Atg1–Atg13–Atg17 complex assembly
Autophagy initiation
Translation, ribosome biogenesis
Cell in state of growth
B
WT
Δmdm1
SCD
SD-N
SCD
SD-N
P-Atg13
Atg13
C
SCD
SD-N
WT
Δmdm1
WT
Δmdm1
P-Sch9
Sch9
Total protein
D
ns
P-Sch9/Sch9 (SCD)
0.3
0.2
0.1
0.0
WT
Δmdm1
E
WT
Δmdm1
SCD
Rap
CHX
SCD
Rap
CHX
P-Sch9
Sch9

### Fig. S3

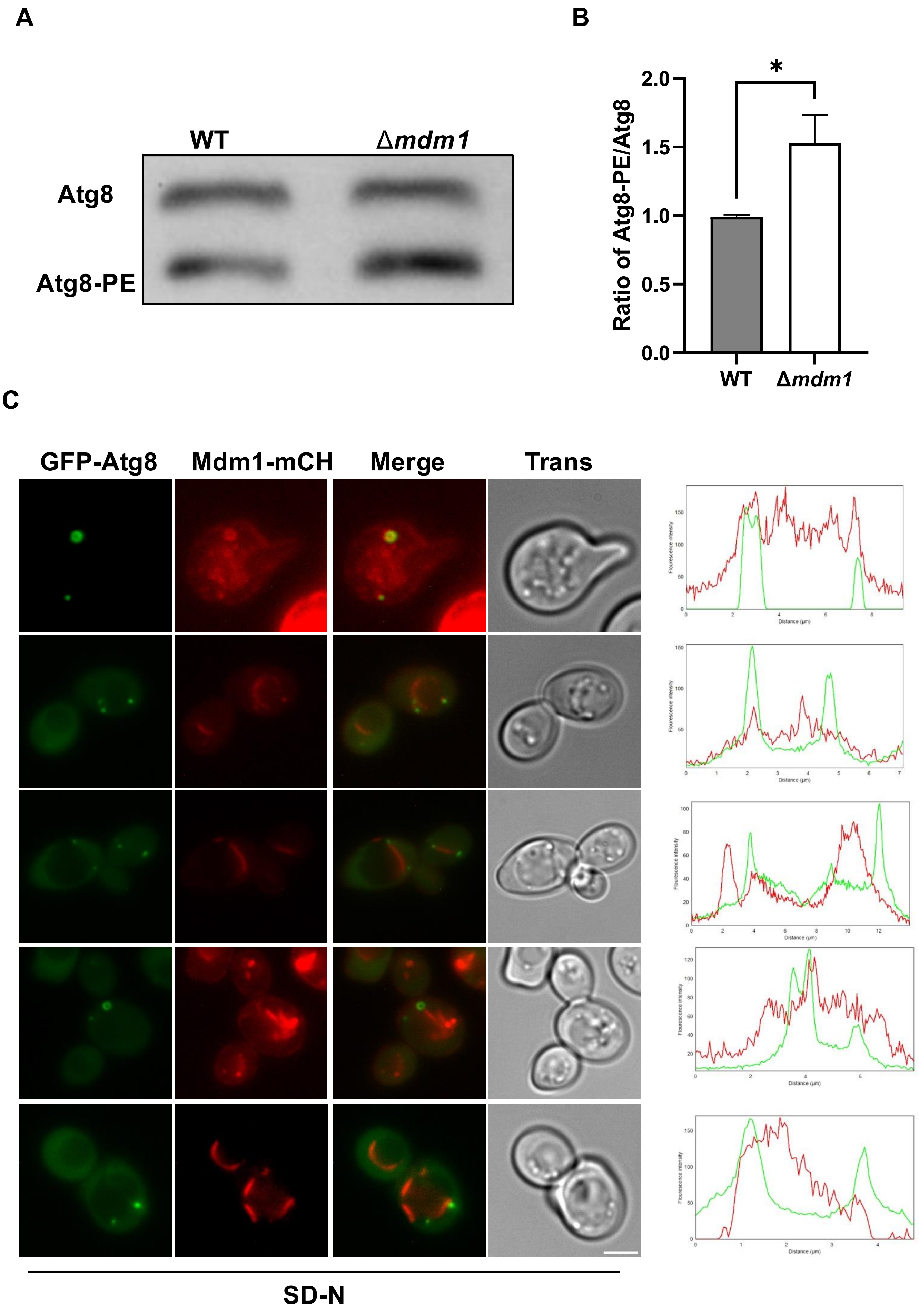
A
WT
Δmdm1
Atg8
Atg8-PE
B
Ratio of Atg8-PE/Atg8
*
2.0
1.5
1.0
0.5
0.0
WT
Δmdm1
C
GFP-Atg8
Mdm1-mCH
Merge
Trans
SD-N

### Fig. S4

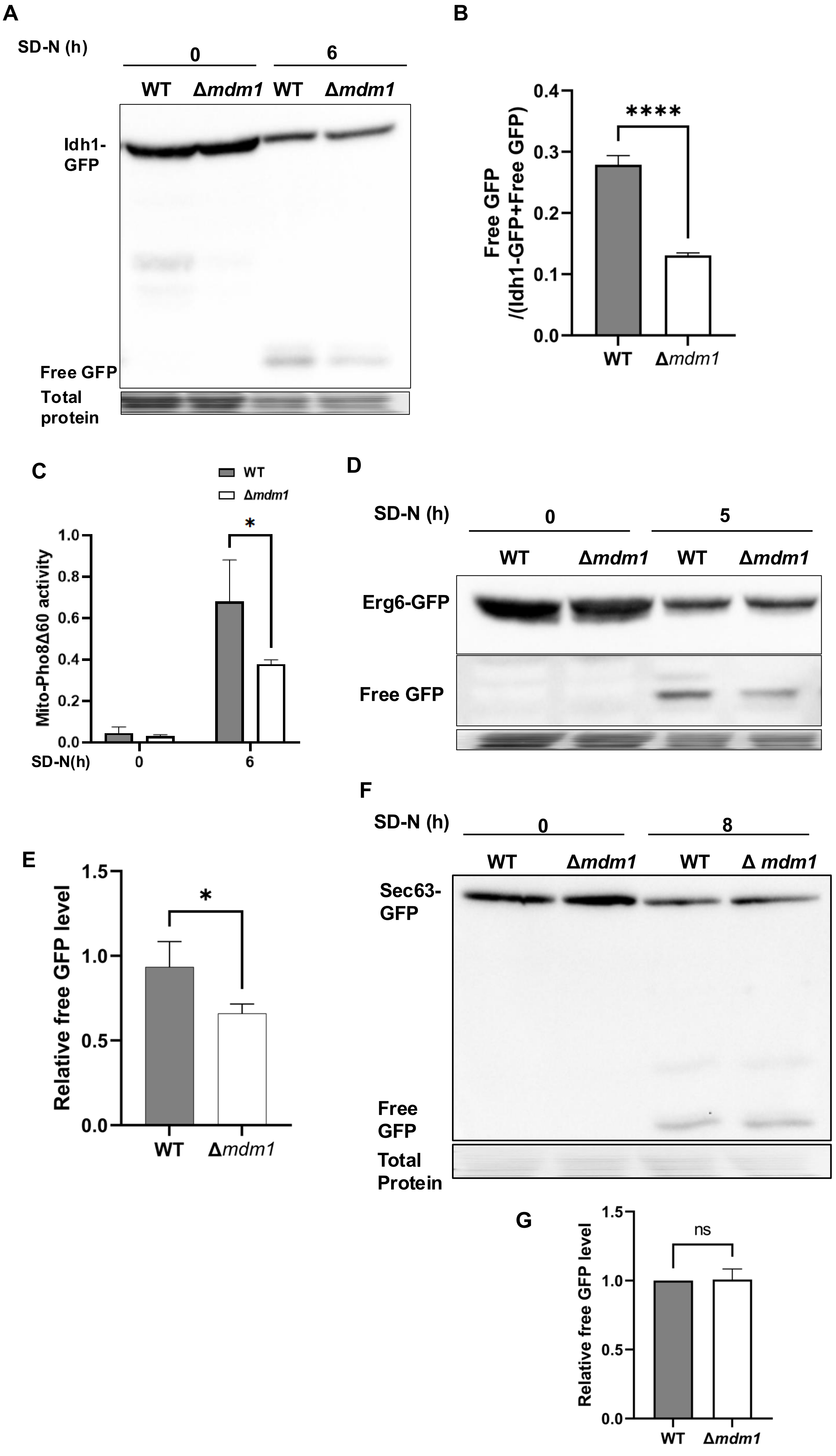
A
SD-N (h)
0
6
WT
Δmdm1
WT
Δmdm1
Idh1-GFP
Free GFP
Total protein
B
Free GFP /(Idh1-GFP+Free GFP)
0.4
0.3
0.2
0.1
0.0
****
WT
Δmdm1
C
WT
Δmdm1
Mito-Pho8Δ60 activity
1.0
0.8
0.6
0.4
0.2
0.0
*
SD-N(h)
0
6
D
SD-N (h)
0
5
WT
Δmdm1
WT
Δmdm1
Erg6-GFP
Free GFP
E
Relative free GFP level
1.5
1.0
0.5
0.0
*
WT
Δmdm1
F
SD-N (h)
0
8
WT
Δmdm1
WT
Δ mdm1
Sec63-GFP
Free GFP
Total Protein
G
Relative free GFP level
1.5
1.0
0.5
0.0
ns
WT
Δmdm1

### Fig. S5

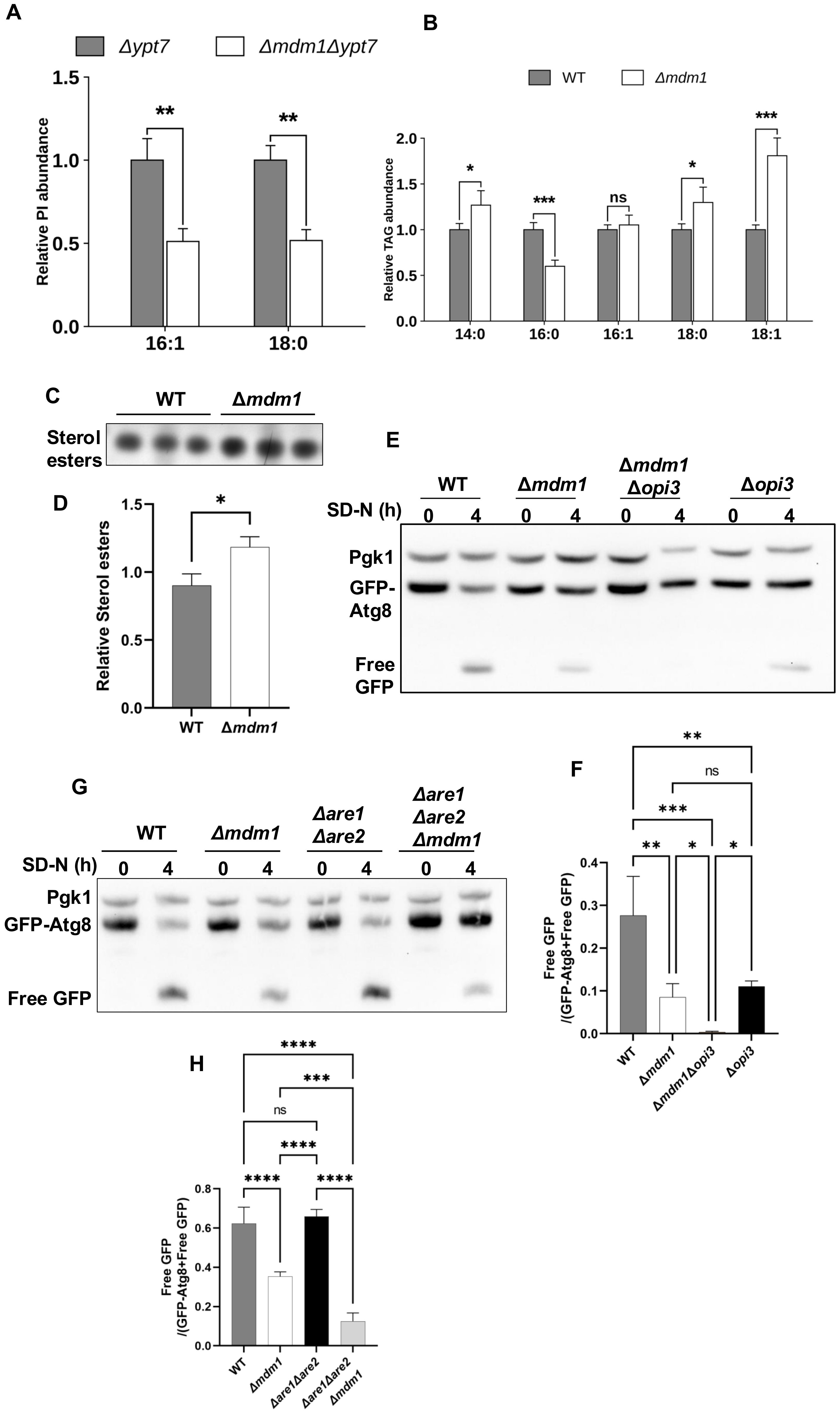
A
Δypt7
Δmdm1Δypt7
Relative PI abundance
1.5
1.0
0.5
0.0
**
**
16:1
18:0
B
WT
Δmdm1
Relative TAG abundance
2.0
1.5
1.0
0.5
0.0
*
***
ns
*
***
14:0
16:0
16:1
18:0
18:1
C
WT
Δmdm1
Sterol esters
D
Relative Sterol esters
1.5
1.0
0.5
0.0
*
WT
Δmdm1
E
WT
Δmdm1
Δmdm1 Δopi3
Δopi3
SD-N (h)
0
4
0
4
0
4
0
4
Pgk1
GFP-Atg8
Free GFP
F
**
ns
***
**
*
*
Free GFP /(GFP-Atg8+Free GFP)
0.4
0.3
0.2
0.1
0.0
WT
Δmdm1
Δmdm1Δopi3
Δopi3
G
WT
Δmdm1
Δare1 Δare2
Δare1 Δare2 Δmdm1
SD-N (h)
0
4
0
4
0
4
0
4
Pgk1
GFP-Atg8
Free GFP
H
****
***
ns
****
****
****
Free GFP /(GFP-Atg8+Free GFP)
0.8
0.6
0.4
0.2
0.0
WT
Δmdm1
Δare1Δare2
Δare1Δare2 Δmdm1

### Fig. S6

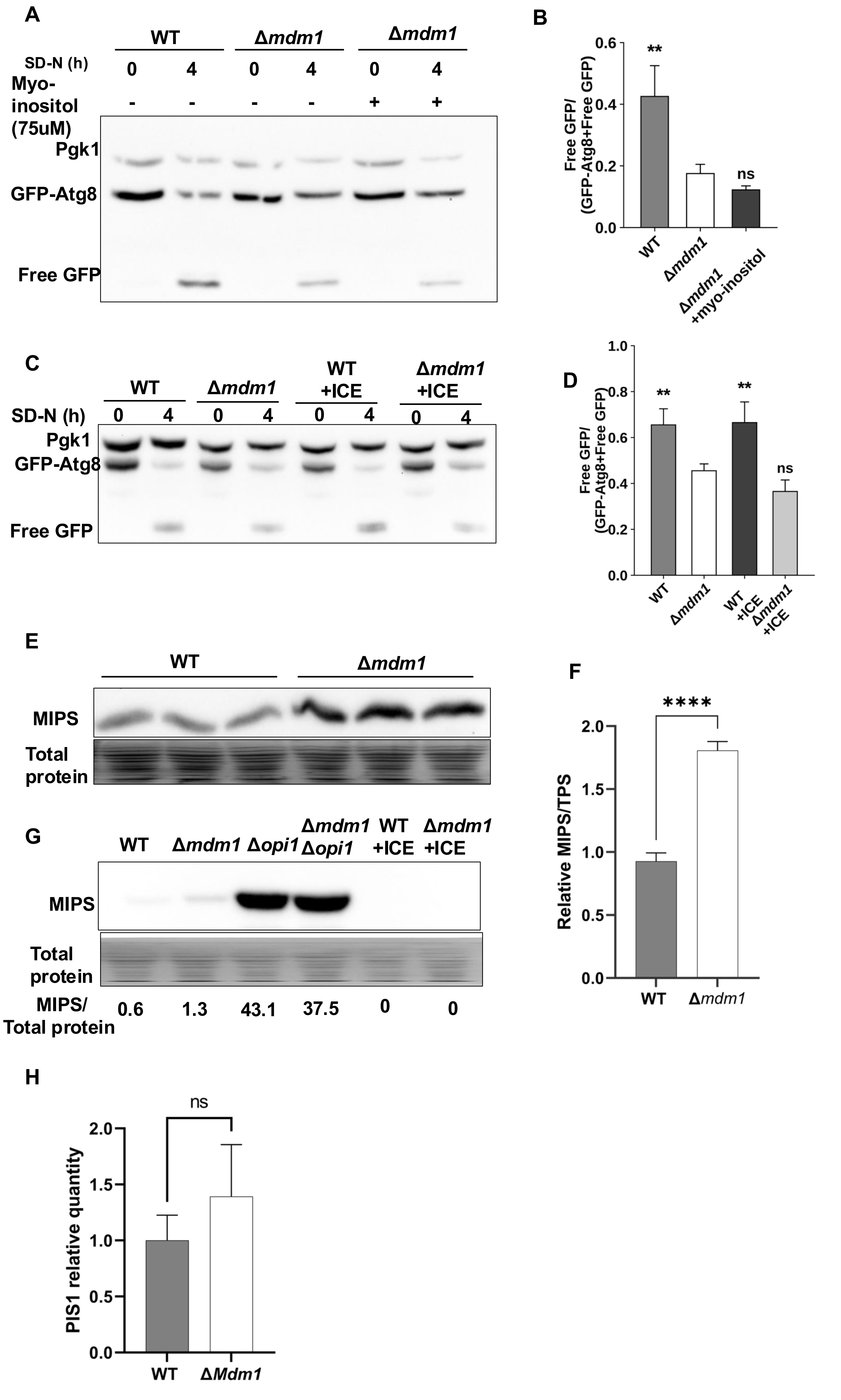
A
WT
Δmdm1
Δmdm1
SD-N (h)
0 4 0 4 0 4
Myo-inositol (75uM)
- - - - + +
Pgk1
GFP-Atg8
Free GFP
B
Free GFP/ (GFP-Atg8+Free GFP)
0.6
0.4
0.2
0.0
**
ns
WT
Δmdm1
Δmdm1 +myo-inositol
C
WT
Δmdm1
WT +ICE
Δmdm1 +ICE
SD-N (h)
0 4 0 4 0 4 0 4
Pgk1
GFP-Atg8
Free GFP
D
Free GFP/ (GFP-Atg8+Free GFP)
1.0
0.8
0.6
0.4
0.2
0.0
**
**
ns
WT
Δmdm1
WT +ICE
Δmdm1 +ICE
E
WT
Δmdm1
MIPS
Total protein
F
****
Relative MIPS/TPS
2.0
1.5
1.0
0.5
0.0
WT
Δmdm1
G
WT
Δmdm1
Δopi1
Δmdm1 Δopi1
WT +ICE
Δmdm1 +ICE
MIPS
Total protein
MIPS/ Total protein
0.6
1.3
43.1
37.5
0
0
H
ns
PIS1 relative quantity
2.0
1.5
1.0
0.5
0.0
WT
ΔMdm1

### Fig. S7

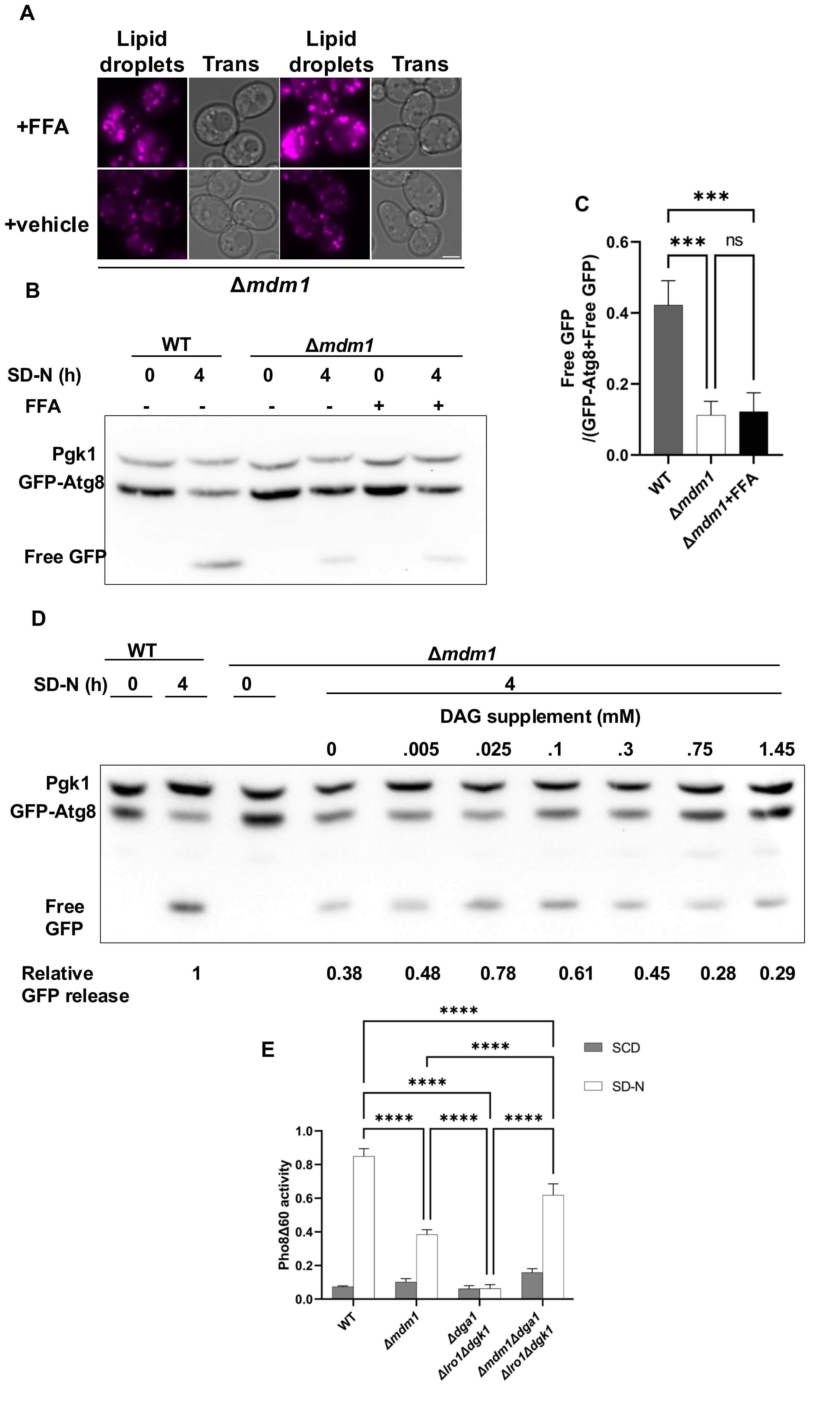
A
Lipid droplets
Trans
Lipid droplets
Trans
+FFA
+vehicle
Δmdm1
B
WT
Δmdm1
SD-N (h) 0 4 0 4 0 4
FFA - - - - + +
Pgk1
GFP-Atg8
Free GFP
C
Free GFP /(GFP-Atg8+Free GFP)
0.0
0.2
0.4
0.6
***
***
ns
WT
Δmdm1
Δmdm1+FFA
D
WT
Δmdm1
SD-N (h) 0 4 0 4
DAG supplement (mM)
0 .005 .025 .1 .3 .75 1.45
Pgk1
GFP-Atg8
Free GFP
Relative GFP release 1 0.38 0.48 0.78 0.61 0.45 0.28 0.29
E
SCD
SD-N
****
Pho8Δ60 activity
0.0
0.2
0.4
0.6
0.8
1.0
WT
Δmdm1
Δdga1 Δlro1Δdgk1
Δmdm1Δdga1 Δlro1Δdgk1

### Fig. S8

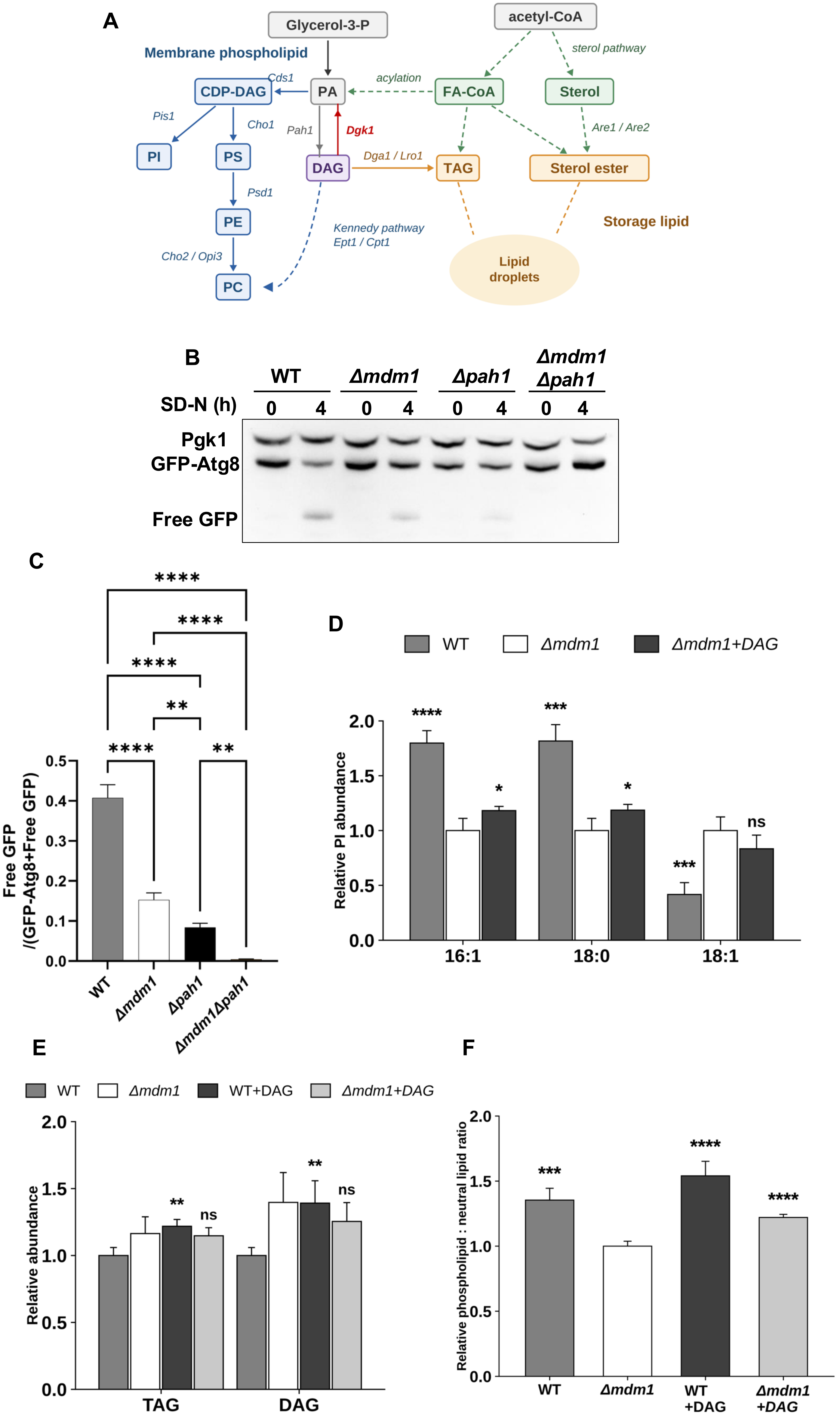

A
Glycerol-3-P
acetyl-CoA
Membrane phospholipid
sterol pathway
Cds1
acylation
CDP-DAG
PA
FA-CoA
Sterol
Pis1
Cho1
Pah1
Dgk1
Are1 / Are2
PI
PS
DAG
Dga1 / Lro1
TAG
Sterol ester
Psd1
PE
Kennedy pathway
Ept1 / Cpt1
Storage lipid
Cho2 / Opi3
Lipid droplets
PC
B
WT
Δmdm1
Δpah1
Δmdm1 Δpah1
SD-N (h)
0 4 0 4 0 4 0 4
Pgk1
GFP-Atg8
Free GFP
C
Free GFP /(GFP-Atg8+Free GFP)
0.5
0.4
0.3
0.2
0.1
0.0
****
****
****
**
****
**
WT
Δmdm1
Δpah1
Δmdm1Δpah1
D
WT
Δmdm1
Δmdm1+DAG
Relative PI abundance
2.0
1.5
1.0
0.5
0.0
****
***
*
*
***
ns
16:1
18:0
18:1
E
WT
Δmdm1
WT+DAG
Δmdm1+DAG
Relative abundance
2.0
1.5
1.0
0.5
0.0
**
ns
**
ns
TAG
DAG
F
Relative phospholipid : neutral lipid ratio
2.0
1.5
1.0
0.5
0.0
***
****
****
WT
Δmdm1
WT +DAG
Δmdm1 +DAG
